# CICLOP: A Robust, Faster, and Accurate Computational Framework for Protein Inner Cavity Detection

**DOI:** 10.1101/2020.11.25.399246

**Authors:** Parth Garg, Sukriti Sacher, Prutyay Gautam, Mrinal, Atul, Arjun Ray

## Abstract

Internal cavities in proteins are of critical functional importance. They can serve as substrate/ligand binding sites, pave path for movement of bio-molecules and even mediate structural conformations occurring between domain interfaces during structural transitions. Yet, there is a paucity of computational tools that can accurately, and reliably characterize the inner cavities of the proteins, a prerequisite for elucidating their functions. We have developed a novel method, CICLOP, that can accurately identify these regions at an atomic resolution. The method is able to accurately detect residues lining the inner cavity, the diameter and volume occupied by the cavity, as well as physicochemical properties of residues lining the cavity such as their hydrophobicity and secondary structure distribution in detail. Additionally, our method also provides an option for computing conservation scores for the residues detected on the inside, allowing for a thorough functional characterization of the cavity.

**Availability:** CICLOP is available at http://ciclop.raylab.iiitd.edu.in/. A compiled Linux executable can be downloaded from https://ciclop.raylab.iiitd.edu.in/standalone/

## 1 Introduction

Proteins are the major drivers of diverse cellular processes. The functionality of these bio-molecular machines largely depends on their three-dimensional structure, accurate estimation of which is an ever-challenging task. For several proteins, presence of functional topological features such as pockets, clefts, tunnels, pores, voids and internal cavities further increase their structural complexities. Amongst these structural features, pockets refer to shallow depressions on the protein surface, while clefts imply a deeper pocket. A channel or a tunnel refers to a path that connects the protein’s exterior with an internally localized point and a pore is a type of channel that traverses the entire molecule. However, internal cavities are a special class of voids that may be open from one or both ends, as in tunnels/channels, or completely enclosed. These features serve as ligand binding sites ^13^, active or allosteric sites in enzymes ^6^, channels for transportation of small molecules ^30^, interfaces for protein oligomerization or for protein-protein interactions ^15;26^ and sometimes just niche environments excluded from the bulk solvent ^28^. The molecular composition and the physicochemical properties of these cavities is cognizant of the underlying protein folding mechanism ^10^ and can impart specificity and selectivity towards their cognate biomolecules ^23^.

In the past decade, several computational methods have been devoted to characterizing tunnels or channels in transmembrane proteins such as PoreWalker ^19^, Hole ^24^, MOLE*online* ^22^, Caver Web ^27^, that utilize various geometry-based algorithms to compute results. However, a vast majority of these are limited by their functionality, accuracy, automation and comprehensiveness, some even require user defined starting points for identification of the tunnel. These approaches limit their usefulness to a select few well-annotated proteins where information about the coordinates lying in either the catalytic pockets, or the largest void enclosed, or channel is available ^27^. Moreover, several proteins have additional pockets or cavities which are secluded from the largest void and characterization of these spaces is overlooked. All the methods that have been developed till date mostly cater towards channels/tunnels that host an entrance/exit. To surpass these limitations, we have developed CICLOP (Characterization of Inner Cavity Lining Of Proteins), an end-to-end automated novel solution for the identification and characterization of protein cavities at an atomic resolution. CICLOP builds on a novel algorithm that imparts unprecedented accuracy, and reproducibility, allowing it to outperform its predecessors. Our method is available both as a web server as well as a standalone, allowing users to perform an in-depth analysis by merely uploading a protein structure file (PDB format).

CICLOP’s algorithm maps the protein structure to a 3D grid consisting of cubes and then performs a breadth-first search to find all the continuous empty regions. Subsequently, by ‘slicing’ the protein along the central pore axis into several thin sections, it elucidates the lining of the cavity. This is further processed to yield a list of residues that face the cavity, diameter and volume profile along the pore axis, secondary structure distribution as well as physiochemical properties of residues that make up the cavity, in terms of their hydrophobicity and charge distribution as one moves along the pore axis. CICLOP can also compute the conservation scores of residues lining the cavity implicating their evolutionary roles.

This study highlights the versatility of our tool in effectively identifying and characterizing cavities of various conformations and sizes that make up functionally diverse proteins. The paper describes the practicality of the various state-of-the-art analytical modules included in our method and their utilization for previously uncharted biological inferences. We have also performed in-silico validation of the results obtained by CICLOP to demonstrate the unparalleled accuracy, precision and sensitivity of our method. Lastly, we have qualitatively and quantitatively compared our algorithm’s performance against other existing popular methods to establish CICLOP’s multifarious utility and superior performance in characterizing protein cavities at an atomistic resolution.

## 2 Approach

CICLOP maps and identifies the different residues that line the inner surface of any cavity-containing protein. Each job on the server mandates the user to provide their email address along with the PDB file(s) of the proteins which the user wishes to characterize.

### Alignment of the Structure

The user is offered two choices for the alignment of their protein structure(s). In the automatic mode, the protein structure is aligned based on the best-fit line that describes the positions of all the alpha carbon atoms. The protein is then rotated such that the best fitting line lies along the positive Z-axis. The protocol of automatic alignment of the input structure has been tested rigorously for reproducibility (deviation from mean of number of residues detected by CICLOP lies within 1% (Supplementary Figure 1). In the manual alignment, CICLOP assumes that the provided structure has the best-fit line (or the central pore axis) aligned with either the negative or the positive Z-axis.

### Identification and Mapping of the Inner Surface Residues

The identification and mapping of inner surface residues by CICLOP is a six stage process.

#### Stage I

All the relevant information (such as coordinates of all atoms, the residues and protein chains) is extracted from the given PDB file. All the atoms of the structure are subjected to linear transformation, which places the protein in the first octant of the three dimensional system with coordinates of all atoms having either a positive value or a zero value.

#### Stage II

The protein after the linear transformation is mapped to a three-dimensional grid of cubes similar to voxels (A voxel is a unit of graphic information that defines a point in three-dimensional space). Each constituent voxel cube has a volume of 1 Å^3^. It may be noted that each cube in the three dimensional grid may either contain no atom, one atom, or more than one atom. The cubes containing no atoms are identified and marked empty. Since each voxel cube from the complete grid contains references to cubes bounding it from all of the six faces, therefore, each voxel also includes information about voxels lying on its boundaries, allowing this data to be represented as a directed graph. Each node defining the positions of a group of atoms lying in close vicinity and each edge defining the relative positions of nodes in the three dimensional space.

#### Stage III

A breadth-first search algorithm is applied to look for empty “nodes” starting from the node containing coordinates of the geometrical center of the transformed protein. If a node is empty, it may be part of a cavity inside the protein. Therefore, edges emanating from this node are traversed. The search continues in all the six cardinal directions until no more empty nodes are encountered. This region physically represents the boundary of the protein (both, the inner and the outer). The atoms surrounding the identified empty regions are initially marked to be part of the inner surface of the protein. This is done to separate them from the bulk of the protein.

#### Stage IV

The next step involves separating the inner surface from the outer surface. The algorithm for this step involves making 1 Å slices of the protein along the Z-axis. For every such slice made:

- The atoms marked to be on the inner surface are identified.
- The mean radius at which these residues lie from the geometrical centre of these marked atoms is identified.

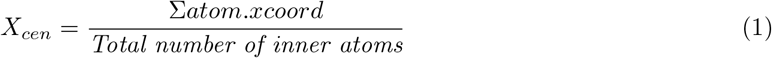

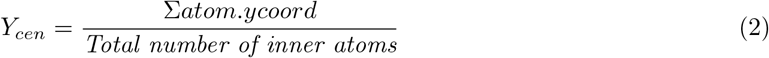

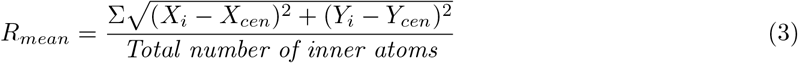 Where *X*_*cen*_ and *Y*_*cen*_ are the X and Y coordinates of the geometric centre of the inner atoms identified for the given Z slice. *R*_*mean*_ is the mean distance at which these inner atoms lie around the given Z coordinate
- The standard deviation of all the atoms (lying at a distance less than the *R*_*mean*_ from the geometrical centre) is calculated from the *R*_*mean*_.

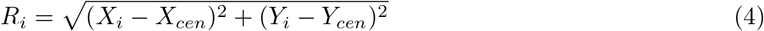

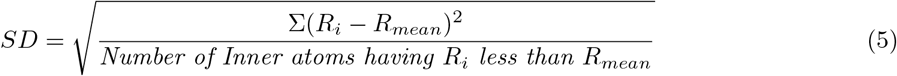 Where *R*_*i*_ is the distance from (*X*_*cen*_, *Y*_*cen*_) along the XY plane, *R*_*mean*_ is the mean radius and *SD* is the Standard Deviation of the *R*_*i*_ from *R*_*mean*_ ∀ atoms having *R*_*i*_ ≤ *R*_*mean*_
- Any atom lying outside the circle of *Radius* = *R*_*mean*_ − (0.7) × *SD* is unmarked and removed from the list of atoms lying on the inside lining. Stages ***II***, ***III*** and ***IV*** are repeated until all the empty cubes have been traversed at least once.

#### Stage V

Once all the atoms lining the inner surface have been identified, a PDB file is rewritten with the original coordinates and a new B-factor value such that the temperature factor 9999 is assigned to atoms lying on the inner surface while a value of 0 is allotted to all others. This ensures that the user can effectively visualize the atoms lying on the inner surface identified by CICLOP.

#### Stage VI

Finally, residues to which these atoms belong are written in a separate file called ’residue.dat’ which is provided as one of the outputs. The list of residues that line the surface contains all those residues for which a single atom was detected to be on the inner surface. Considering these calculations are performed on static structures, even if a single atom of a residue is detected on the inner lining, under real-life dynamic situations, it is likely that the complete residue becomes available for interactions in the cavity. A schematic representation of the algorithm is depicted in Supplementary Figure 2.

### Calculation of the pore diameter profile

The calculation of the pore diameter is based on the largest circle that can fit inside the pore at the given length along the pore axis. The diameter measurements were benchmarked to be recorded for every 3 Å block. This cut-off was chosen because most of the atoms of a protein have a mean van der Waals diameter of ~ 3.6 Å ^20^, and therefore, the diameter is unlikely to change much at a smaller step size (Supplementary Figure 3). The pore diameter is calculated by calculating the Voronoi diagram of all the atoms that are identified on the inner surface in the 3 Å block in order to obtain its Voronoi vertices.

If *X* is a metric space with a distance function *d*. Let *K* be a set of indices and (*P*_*k*_)_*k*∈*K*_ be an ordered collection of non-empty sites in space *X*. The Voronoi Region (*R*_*k*_), associated with the site *P*_*k*_ is the set of all points in *X* whose distance to *P*_*k*_ is not greater than their distance to other sites in *X*.

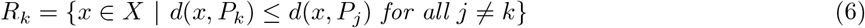

Voronoi vertices are the points where three or more of the Vornoi regions (*R*_*k*_) intersect. The distance used in our study is the familiar *Euclidean distance* that is calculated as follows:

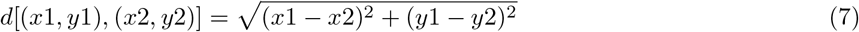

Similarly, a convex hull is calculated for all the atoms lying on the inner surface in the same block. For all Voronoi vertices lying inside the convex hull, the distance between the inner surface atom closest to the Voronoi vertex is marked as a possible radius. The maximum value of all such radii is doubled and 3 Å is subtracted from that value to account for van der Waals radii from the opposite ends. The value so obtained is the diameter for the given block. This process is repeated for all the 3 Å blocks and plotted as a diameter along the Z-axis plot, which is provided as an output to the user.

### Calculation of the pore volume profile

The pore volume is calculated for every 3 Å block along the pore axis.The total volume of a block is the sum of the volumes of the three individual 1 Å slices cut perpendicular to the pore axis that makes up the block. For every 1 Å slice, its pore volume value is obtained by first calculating the geometric centre of all the atoms lining the inner surface of that slice. Taking the centre as the reference point, this list of atoms is sorted in a clockwise direction. On joining every unique pair of consecutive points with the centre, a triangle is formed, and the area of all such triangles is calculated and summed. The division of the entire circular slice into smaller triangles that all have the centre of the circle as a common vertex ensures that no two triangles have overlapping areas. The slice volume is measured by multiplying the height of the slice (1 Å in this case) with the total area of the slice.

The area for each triangular region is calculated using the widely used *Heron′s Formula*.

## 3 Methods

### Visualization of Cavity Volume

A PQR format file, provided along with the results, can be used for visualizing the cavity. The file contains information regarding the coordinates where spheres of 1 Å radius are constructed such that they occupy the entire space of the detected cavity at different points along the pore axis. For every 1 Å step along the pore axis, the circles formed according to the radii are filled with *n* spheres placed uniformly inside the circles. The number of spheres is a function of the cavity radius at the point and is given by the following formula:

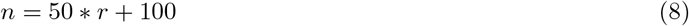

where r is the radius of the cavity along the pore axis and n is the number of spheres used to cover the cavity at the Z coordinate.

### Creation of Hydrophobicity Distribution Plot

To create this plot, CICLOP traverses the identified pore once again along with the identified (or provided) pore axis. For each Å step, the residues lining the pore are taken into consideration. The summation of their hydrophobicities as indicated on the Kyte-Doolittle scale are plotted as a function of the distance along the central axis of the pore.

### Creation of Charge Distribution Plot

Similar to the previous plot, the central pore is traversed along the central axis. For each Å step, the total number of positively and negatively charged amino acids is plotted as a function of the distance along the pore axis. Amino acids LYS, ARG, and HIS are taken as positively charged, while ASP and GLU are considered to be negatively charged.

### Evaluating Residue Conservation and Calculation of the Conservation Scores

Extracting the sequence from the structure, basic local alignment search is performed using local nr database. Post generating the multiple sequence alignment file, evolutionary scores are calculated and mapped onto the structure (Supplementary Figure 4).

### Collection of Data Used in This Study

All the protein structures used in this study were obtained from the RCSB database, and their PDB IDs have been indicated in their respective figures. The structures used in the ABCA1 case study was 5XJY while those in ATP synthase case study was as follows: 5ARA (State 1A), 5ARE (State 1B), 5ARH (State 2A), 5ARI (State 2B), 5FIJ (State 2C), 5FIK (State 3A), 5FIL (State 3B). All the images used in this study were produced using UCSF Chimera ^21^ (except the full structure of ATP synthase, which was produced using Chimera X ^8^. In the ABCA1 case study, the list of reported SNPs was obtained from dbSNP using a ”pathogenic and likely pathogenic” filter.)

### Identification of Inner Residues Using Various Tools

For PoreWalker, each structure was uploaded on their web server, and the PDB file containing the marked inner atoms was downloaded. All the residues were extracted from the B-factor loaded PDB using an *in-house* Python program.

For MOLE*online*, each structure was submitted to their web server with the default parameters. Option to “Ignore HETATOMs” was selected. The options “Merge Pores” and “Automatic Pores” were turned on. Once the calculations were over, the results were downloaded and the inner residues extracted using an *in-house* Python script.

For Caver Web, after submitting the structure, the catalytic pocket identified by the tool with the highest pocket score was taken as the starting point for the calculations. In case Caver Web was unable to identify a starting point, some inner surface residues were selected manually as indicated in the literature. Upon downloading the results, the inner lying residues were again extracted using a Python protocol developed for this purpose.

For CICLOP, all the structures were submitted in the automatic mode (except PDB IDs 1LNQ and 2OAR) of alignment, unless stated otherwise. In cases where conservation scores were calculated, the underlying evolutionary model was selected as ‘LG’ unless otherwise stated. All the conservation scores calculated for the purpose of this study were done through the empirical bayesian methodology with no exceptions.

### Molecular Dynamics Simulation for Quantitative Comparison

All-atom MD simulations were performed using GROMACS ^1;14^, version 2020. OPLS-AA/L all-atom force field ^12^ was used to describe the system. SPC/E water model ^16^ was used to solvate the protein, and sodium ions were added to neutralize the system. All the simulations were performed at 298K using the modified Berendsen thermostat ^3^ for temperature control and Verlet cut off scheme for searching the neighbouring grid cells. Pressure coupling (coupling time 2.0 ps, isothermal compressibility 4.5e-5) using the Parrinello-Rahman scheme was also used in all the simulations due to which the lateral and the perpendicular pressures were coupled independently to maintain a constant pressure of 1 bar. The simulations were performed under periodic boundary conditions in all the cardinal directions, with the long-range electrostatic interactions being treated with the Particle Mesh Ewald (PME) method using a grid-spacing of 0.16 nm combined with fourth-order B-Spline interpolation to compute the potential and forces in between grid points. LINCS algorithm was used to constrain all the bonds. The short-range interactions were cut-off at 1.0 nm. The production run for each system was performed for a total of 10 ns, with two fs being the time step used for numerical integration of the equations of motion. All the starting structures were subjected to a minimization protocol of 50000 steps using the steepest descent algorithm followed by equilibration runs in NVT and then NPT ensembles for 100 ps each.

Each frame post-stabilisation of the protein backbone thus generated by the MD protocol described above was analysed separately using *in-house* Python protocols. Initial frames were discarded in order to account for short fluctuations at the beginning of the simulation. First, all the water atoms lying inside the protein cavity were identified. The distances for each protein atom from the water group identified previously were calculated using GROMACS trjorder. All the residues having at least one atom in close proximity (less than or equal to 3.5 Å) to the water group lying inside the cavity were identified as inner residues for the frame in question. Finally, only those residues were considered to be truly inside the cavity, which appeared to be lying on the inner surface in at least 90% of the frames. The final list of residues thus generated was taken as the “True Positive” set of the inside lying residues, while the rest were categorised as the “True Negatives” for the experiment. Using the residue lists curated in the aforementioned step as true negatives and positives, accuracy and precision was calculated for CICLOP, PoreWalker, MOLE*online* and Caver Web.

### Analysis of ATP Synthase Structures Using CICLOP

#### Alignment of the Structures

The bovine ATP synthase F1 domain cavity consisting of *α*_3_*β*_3_ chains was isolated from the complete crystal structure using UCSF chimera for each of the seven different substates available as crystal structures. The cavity for state 1A was aligned with the Z-axis manually, and the cavities for the rest of the six states were superimposed on the manually aligned structure of state 1A to keep the human error to the minimum. All the seven cavities were then submitted to CICLOP keeping the “Mode of Alignment” as “Manual”.

#### Annotation of the Size of the Cavity

The distances to opposite atoms on chains A and F of the F1 domain of bovine ATP synthase, state 1A, (PDBID: 5ARA) were calculated and marked using the distance module of UCSF Chimera. The corresponding points on the diameter profile generated by CICLOP were also annotated.

#### Evaluating the Orientation of *γ* subunit During the Rotary Cycle

In order to evaluate the orientation of the *γ* subunit with respect to the cavity (*α*_3_*β*_3_), each structure was aligned with respect to chain A. Similarly, to evaluate the orientation of the cavity with respect to *γ*, each structure was aligned with respect to the *γ* subunit (chain G).

#### Calculation of Minimum Distance of *γ* Subunit from Each Chain Constituting the Cavity

Moving from the bottom to the top along the Z-axis, 1 Å thick slices were cut. Any slice which did not contain at least one atom from both the cavity (all its constituent chains) and the *γ* subunit was ignored. In each slice, the distances of all the *γ* atoms were calculated from all the atoms corresponding to chains A, B, C, D, E and F. The minimum value obtained for each combination was recorded. A similar calculation was performed for each slice and was plotted.

#### Calculation of the Dynamic as well as the Constant Residues During State Transitions

Residues that were common in all the seven substates were calculated from the list of residues lining the inner cavity provided by CICLOP. An *in-house* Python script was used to calculate the intersection of the seven sets so formed (each set constituting the list of residues detected by CICLOP for that substate). To calculate the residues that were dynamic during each state transition, similarly, the residue list provided by CICLOP was used. Considering each list of residues as a set, the difference between two sets (for states that appear one after the other) was calculated using an *in-house* Python script.

## 4 Results

### Overview of CICLOP

CICLOP is a computational framework that accurately identifies residues lining the inner cavity of proteins. Using the input of the PDB three-dimensional file format, the method operates in two modes of alignment of the protein, normal to the cavity axis. In the automatic mode, the algorithm rotates the input structure such that its central pore axis lies along the Z-axis while in the case of manual mode, the same is assumed (Supplementary Figure 2). The output resultant B-factor loaded PDB output file marks the detected residues forming the inner surface of the cavity. This is followed by estimating the total pore diameter and volume, which is calculated using the sum of the areas enclosed by all the inner lining atoms. Both the diameter and volume are plotted along the length of the pore and are provided to the user (Supplementary Figure 5). Moreover, the user is also provided with a PQR formatted file that can be used to visualize the volume occupied by the internal cavity in three-dimensional space. CICLOP has been tested on a large subset of cavity containing proteins varying in sizes and shape. Supplementary Figure 6 showcases these varied representative protein structures’ internal cavities, where the cavity residues detected by our tool are coloured in red. The heterogeneity in size (174 aa in 2OAR to 994 aa in 1SU4), function (membrane-embedded channel proteins to cytosolic enzymes) that is indicative of the diversity in cavity conformations inherent in biological systems, was exemplified and deciphered using CICLOP.

### Protein cavity analysis using CICLOP

In order to comprehensively annotate protein cavities, CICLOP provides an exhaustive package of easy-to-use features and modules. Our tool can characterize internal cavities at both residue as well as atomistic resolution (Supplementary Figure 7).

In addition to the detection of cavities (Figure 1a), CICLOP computes conservation attributes of cavity-lining residues, which are normalized for comparison between proteins (Figure 1b). Furthermore, the distribution of inner residues based on their conservation score as a function of the cavity length along the axis of the detected cavity is also provided (Figure 1c, Supplementary Figure 8). Additionally, CICLOP assigns the secondary structure for each residue lining the cavity(Figure 1c).

**Figure 1:**
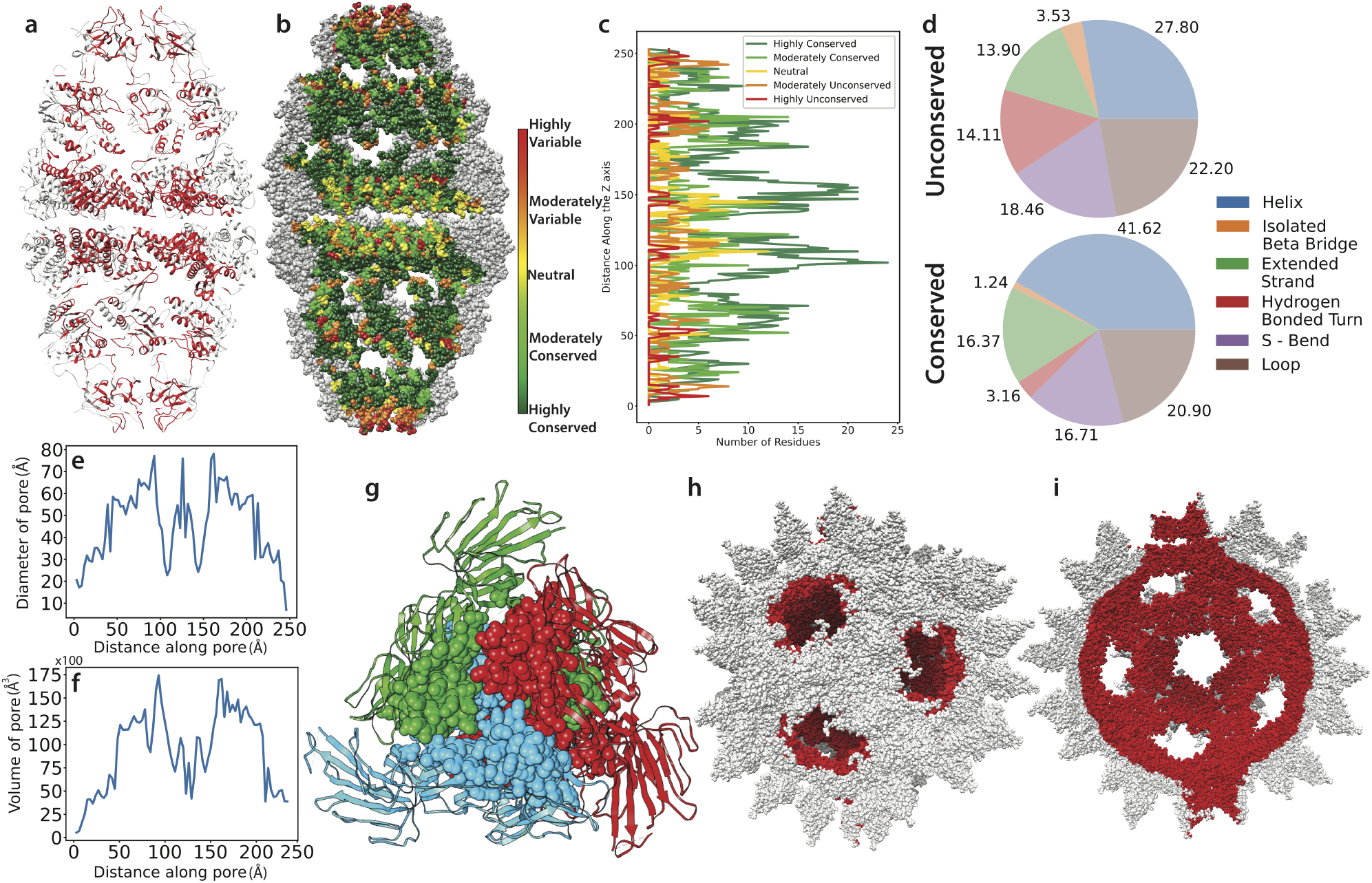
Application of CICLOP. **a.** Identification of the residues lining the inner surface of WT human mitochondrial chaperonin (ADP:BeF3)14 complex (PDB ID: 6HT7). **b.** Conservation of the identified residues; marked in the range of highly variable (red) to highly conserved (dark green)57.83/16.59% of the residues lining the cavity are highly/moderately conserved. **c.** Conservation profile as a function of Z-axis **d.** Secondary structure assignment of the conserved and unconserved residues as detected by CICLOP. **e.** Diameter profile of the pore of the protein as a function of Z-axis distance. **f.** Volume profile of 6HT7 as a function of Z-axis distance (Total pore volume = 738,319.197 Å^3^). **g.** Top view of the oligomerization interface residues (forming a cavity), of the three chains (highlighted in red, blue and green) of the alpha-coronavirus spike glycoprotein (PDB ID: 6IXA), as detected by CICLOP and represented as spheres. **h.** Outer surface of Human parechovirus (HPeV) protein complex (PDB ID: 4UDF) with inner surface marked red and **i.** longitudinal section of the inner surface (red) as detected by CICLOP.

Our method also generates a diameter and volume profile of the cavity detected in the input structure. These profiles are provided along with the output summary file. Figure 1e,f & Supplementary Figure 5 show the diameter and volume profile of the multi-subunit chaperone complexes human and bacterial in origin, respectively. Additionally, CICLOP also facilitates functional characterization of the cavity by providing a charge distribution analysis as well as a hydrophobicity plot along the length of the pore.

CICLOP can also detect cavities at the interface of multimers, such as those formed by the homo-trimeric arrangement in the spike protein of SARS-CoV2 virus (Figure 1g), consequently allowing for the identification of oligomerization domains of these proteins. The robustness of CICLOP is highlighted by its ability to effortlessly characterize proteins as large as human parechovirus envelope (HPeV) containing 302,100 atoms arising from 38,580 residues (Figure 1h & i).

### Validation of inner cavity characterization provided by CICLOP

To substantiate the results obtained by various modules of CICLOP, we performed several in-silico validations. Firstly, to verify the validity of residues detected on the cavity’s inner surface by CICLOP, we performed an all-atomistic molecular dynamics simulation. Several protein-water systems were set up and simulated for 10 ns at room temperature (298K) in order to saturate contacts between protein and water(see Methods and Supplementary Figure 9-11). Simulations were performed on proteins varying in size, cavity conformations, and physiological functions, and our tool consistently performed well, detecting inner residues with a precision of 90.85%, 99.15% & 90.01% and an accuracy of 91.52%, 85.22% & 89.07% respectively.

Next, to validate the diameter profile generated by CICLOP, we characterized a unique cavity, that of the F1 domain of the bovine mitochondrial ATP synthase. ATP synthases consist of two functional domains. The F1 domain is formed by a hexameric cap that fits onto a rotor ^11^. We extracted (from the crystal structure) the measurement of the distance between pairs of atoms of the two chains (that are roughly diametrically opposite to each other), which encompasses the ‘cap’ (Supplementary Figure 12a), and compared it with the diameter profile created by our tool. Supplementary Figure 12b illustrates the high-resolution diameter profile generated by CICLOP. The diameter for each slice obtained by CICLOP relative to the cavity’s physical measurement falls within the error limits (mean error = 0.053). Similarly, Supplementary Figure 12c & d depict the length and width of the cavity inferred from the diameter profile. These values corroborate well with the physical measurements performed on the structure and explain the rotor’s tight fit inside the cavity. Together these results highlight the sensitivity of diameter measurements computed by our tool and how they precisely match the physical measurements of the protein structure obtained from the crystal structure.

### Benchmarking CICLOP against other existing tools

Existing methods for identifying protein cavities/channels employ a wide range of algorithms, such as the Monte Carlo approach by HOLE ^25^, grid-based approach by Caver Web ^27;18^, 3D Voronoi diagrams by MOLE *online* ^22;4^ etc. Most of these popular methods require a certain level of user intervention, such as specifying a point or residue(s), which lie on the protein’s inside to be characterized. While the proteins that form channels might be well-annotated, as they contain two visible openings that connect different cellular environments, an internal cavity may be formed by a completely bound void with no, one or multiple openings. Moreover, a protein might contain several internal cavities, and knowledge of residues/points lying in all such regions might not always be available. Figure 2 summarizes the various features and strengths offered by CICLOP in comparison with the other popular methods.

**Figure 2:**
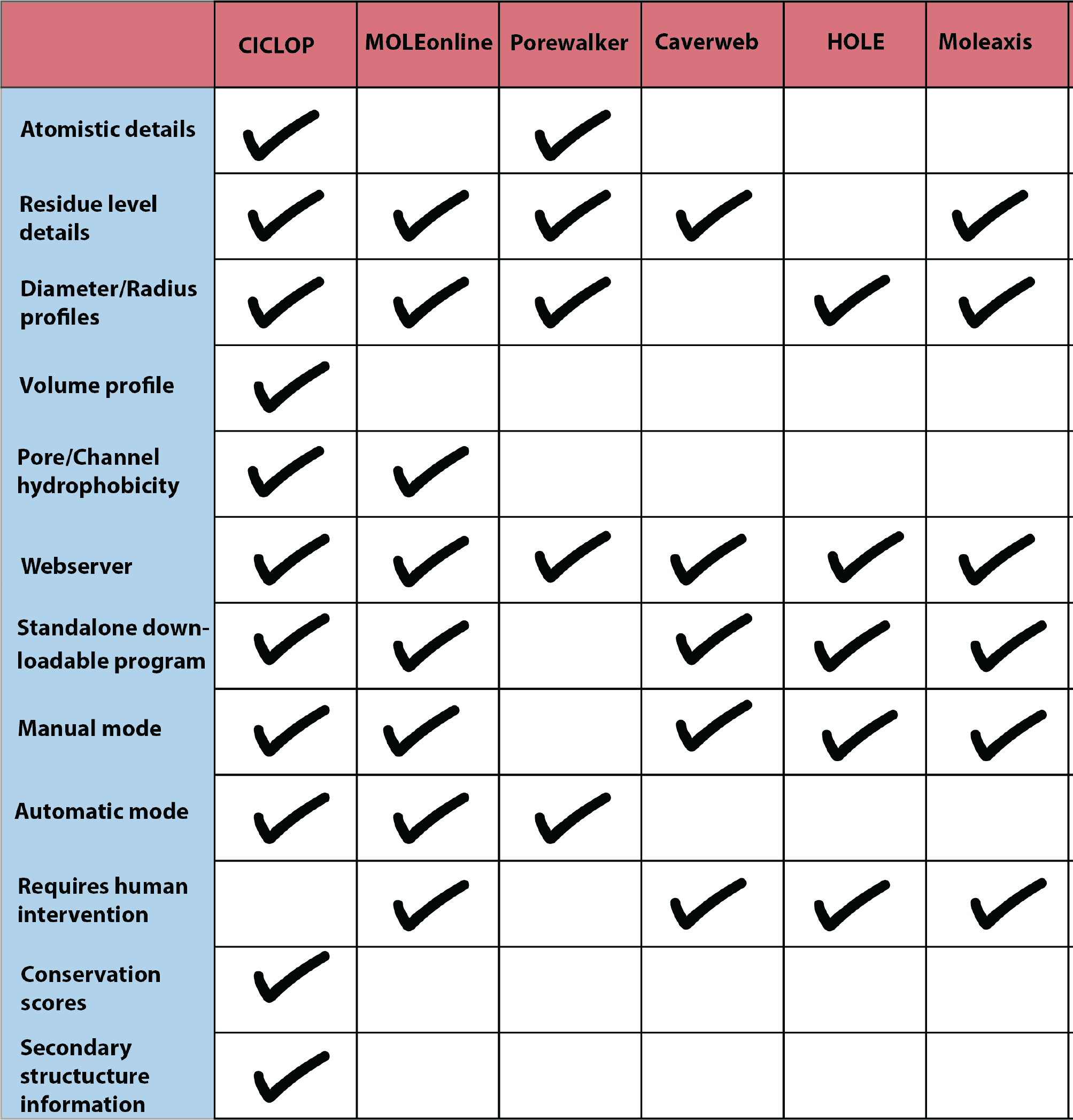
A comparison table of the various features offered by different cavity/tunnel detection tools.

Next, we set out to evaluate CICLOP’s performance against two of the aforementioned methods – PoreWalker, MOLE*online*. We chose a set of 4 proteins [a cytosolic chaperone (1AON), a group II chaperonin (3LOS), Nicotinic AcH receptor (2BG9), and Alpha-hemolysin (7AHL)] from different organisms varying in size, sub-cellular localization, cavity conformation as well as function and used them as input for CICLOP and the other methods. Using only the default parameters for each tool, the inner residues for each input protein were obtained. These were then mapped onto the protein structure (coloured red) and have been displayed in Figure 3. Residues detected by CICLOP form a continuous contour along the cavity and are inline with the protein’s overall structure compared to the other methods (Figure 3a & b). Specifically, MOLE*online* was unable to detect the majority of residues lining the cavity (Supplementary Figure 13 & Figure 3c), and both PoreWalker as well MOLE*online* showed significant ‘leakage’ (defined as residues visibly on the outer surface but detected to be on the inner surface by a method) (Figure 3c & d). Additionally, CICLOP was efficiently able to detect the cavity of a completely closed chaperonin (3LOS) while detections by both PoreWalker as well as MOLE*online* were misplaced. CICLOP’s performance against Porewalker, MOLE*online* and Caver Web was similarly evaluated on 26 additional proteins varying in size as well as cavity conformation (Supplementary Figure 14 & Supplementary Table 1)

**Figure 3:**
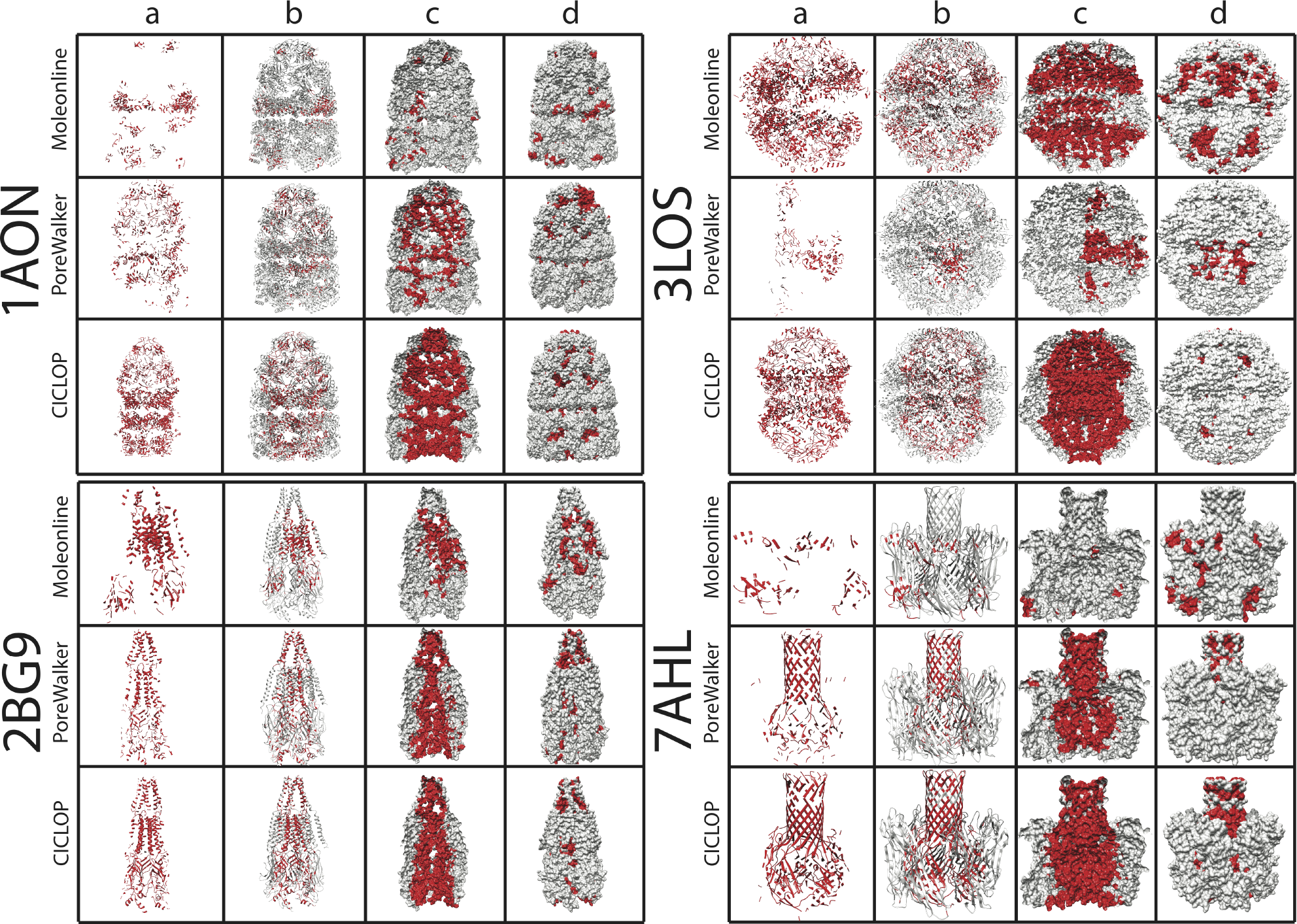
Qualitative comparison of detection of inner-lining residues within a protein cavity with other methods. The residues identified by various methods are marked in red. **a.** The inner surface contour as detected by the different methods. **b.** The inner surface marked (red) for the protein in ribbon representation for different methods. **c.** A longitudinal section of the inner surface (red) of the proteins in surface representation for different methods. **d.** The complete surface as seen from the outside for the corresponding figure in panel c for different methods.

To quantitatively measure each method’s detecting efficiency, we extended the list of true positives obtained from our all-atomistic molecular dynamic simulations to these methods as well. The list of residues lining the cavity obtained by PoreWalker, MOLE*online* as well as Caver Web was similarly compared to the true positive set for each of the three protein structures (1TF7, 6V0B and 1AON), after which the accuracy and precision of detection were computed. PoreWalker gave an accuracy ranging between 62.48% - 70.04% and precision ranging between 72.3% - 98.36%. MOLE*online* performed similarly, giving an accuracy between 58.71% - 67.79% and precision between 31.25% - 77.68%. Surprisingly, Caver Web was unable to process the structure of one of the input structures (1AON) and gave an accuracy in the range 50.13% – 58.74% and precision of 44.82% - 70.79%. Supplementary Figure 9d, 10d & 11d summarize results of CICLOP as well as the three methods. These results highlight that the performance of each of these methods fluctuates significantly with the input structures and hence the size, structure, and cavity conformation of the input protein. While a certain amount of structure-specific performance can be anticipated for each method, CICLOP is significantly consistent in the magnitude of its detection accuracy (85.22% - 91.52%). and precision (90.01% - 99.15%).

Together these results indicate that CICLOP allows for an in-depth characterization of protein cavities varying in size and conformation. Our tool is built on a novel algorithm that imparts it with unprecedented accuracy, and reproducibility such that it outperforms its predecessors.

### CICLOP’s utility in characterizing protein cavities

To demonstrate CICLOP’s applicability and usability in the functional characterization of protein cavities, we used the cavity of the F1 domain of bovine mitochondrial ATP synthase as a case study (Figure 4a). ATP synthases are found in the inner membrane of mitochondria and operate by a rotary catalytic mechanism. Using Cryo-EM, Zhou et al. obtained three rotational states of bovine ATP synthase related to each other by a rotation of 120°. Each of these states was further divided into seven substates providing a snapshot of ATP synthase during its full catalytic cycle ^29^. Investigating the minute changes occurring in the cavity as the central rotor rotated about its axis, we observed that the movement of the central rotor brought about changes in the volume of the cavity (Figure 4b). The overall hydrophobicity of the cavity also followed a cyclic pattern (Figure 4c), increasing from state 1A to 1B and then decreasing during substate transitions of state 2 and then increasing again in state 3A before it reset (state 3B to 1A) The detection sensitivity of our tool is also evident in the comparison of the diameter profiles of state 1A, 2A and 3A generated by CICLOP (Figure 4d-f & Supplementary Figure 15). The changes in the diameter of the top, as well as the bottom part of the cavity (depicted by the two peaks in the profile), also corroborate well with the volume profile of the substates (Figure 4c), implying that the movement of the rotor results in localized conformational changes that change the overall morphometry of the enclosed pore.

**Figure 4:**
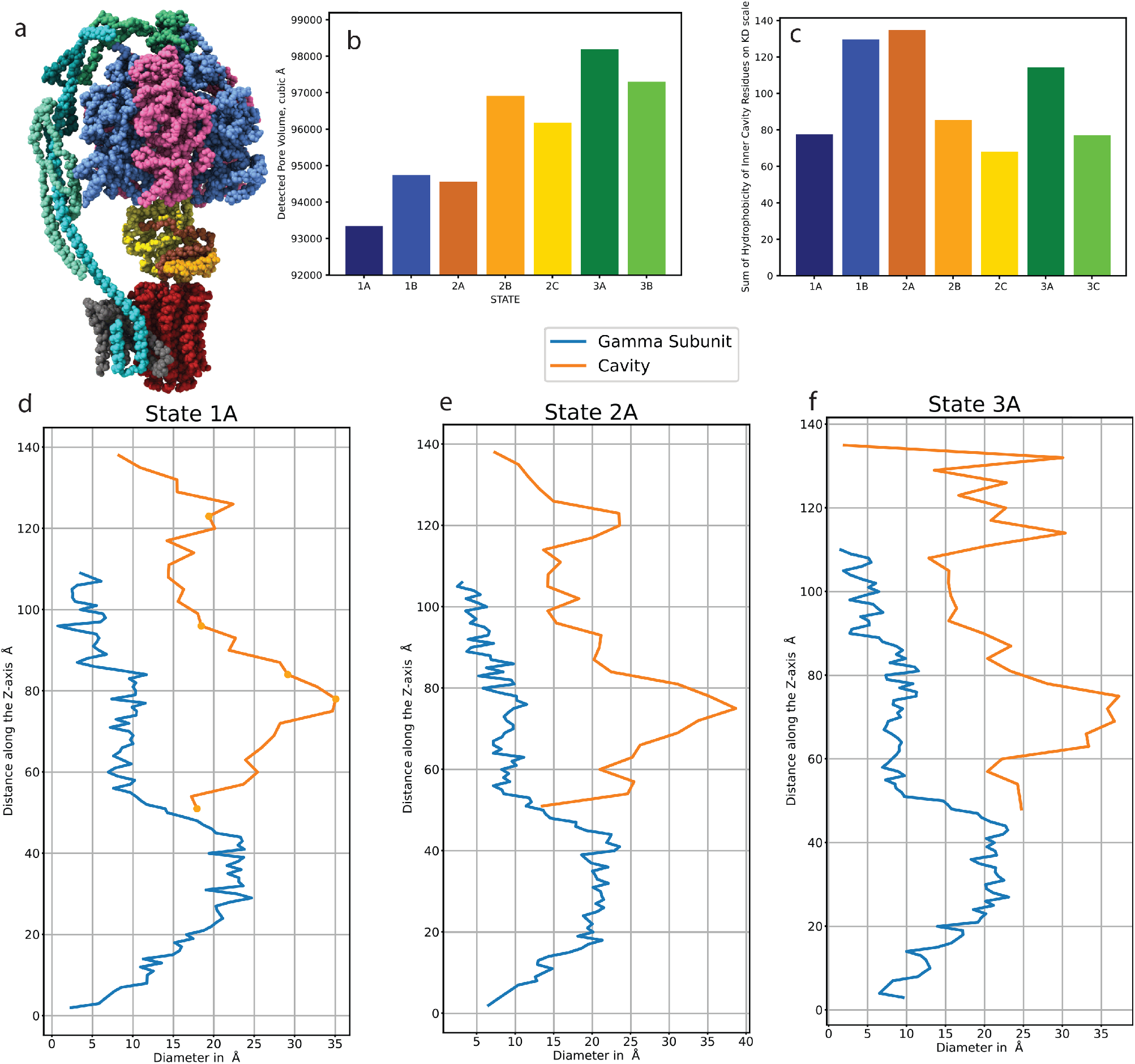
Analysis of the internal cavity of ATP synthase F1 domain during its catalytic rotary cycle using CICLOP. **a.** Structure of bovine mitochondrial ATP synthase (PDBID: 5ARA). Chains A-C(blue) that constitute the *α* subunits as well as chains D-F(pink) that constitute the *β* subunits are alternatively arranged to form the F1 cavity. Chain G (yellow), H (orange) and I (brown) that form the subunit *γ*, *δ*, *ϵ* respectively, together constitute the rotor complex of the F1 domain. Chains J-Q(red) that form the c1 subunit complex of F0 domain are inserted in the mitochondrial membrane. a1 subunit formed by chain W (gray) is attached to c1. The peripheral stalk formed by OSCP complex (chain S), subunit b1 (chain T), subunit d (chain U), subunit f6 (chain V) are coloured as sea green, turquoise, bright blue and teal respectively. **b.** The pore volume of ATP synthase cavity during a complete rotation. **c.** The overall hydrophobicity of the ATP synthase cavity during its rotation. Each state is represented by a differently coloured bar. **d.** The diameter profile generated using CICLOP of the ATP synthase cavity for State 1A. **e.** State 2A **f.** State 3A.

During the rotary cycle of ATP synthase, the F1 domain is speculated to be mostly immobile (albeit a slight bobbing movement due to the torque generated by turning of *γ* subunit) and held in place by a peripheral stalk that connects it to the membrane-embedded region of F0 (Figure 4a) ^9^. As the *γ* subunit complex rotates inside the cavity, it orients itself towards the interface of an *α β* subunit during each 120° transition (Supplementary Figure 16). This also results in conformational changes in the cavity’s external face that *γ* orients towards, characteristic of the nucleotide-binding states (Open, Tight, and Loose) ^2^. We suspected that the transition between the different nucleotide-binding states, although on the outer surface, would be reflected in the cavity interface, as they are caused by the spinning of the rotor inside. This is evident from the changes in cavity composition summarized in Supplementary Figure 17a. Each 120° rotation (state transition) leads to about 15% change in residues lining the cavity. However, these changes are minor during substate transitions. The final transition representing the completion of one entire cycle (from state 3B to 1A) results in tremendous fluctuations in the cavity as computed by CICLOP (30.35% change in residue composition). We expanded on our observation by visualizing the B-factor loaded output generated by CICLOP to deduce the spatial location of residues lining the cavity during substate transitions. Supplementary Figure 17b & Supplementary Figure 18 show that the residues immediately surrounding the *γ* subunit complex are well represented in all the substate transitions. However, there are newer clusters that lie concentric to these, which are now exposed to the cavity, possibly eliciting conformational changes on the outer surfaces allowing cyclic binding and release of nucleotides. Supplementary Figure 17c & Supplementary Figure 19 zooms in on this observation and depicts the major changes in these ‘dynamic residues’ occurring in state transitions versus minor fluctuations evident in sub state transitions. Additionally, we also employed the conservation and secondary structure module included in CICLOP and observed that the internal organization of the cavity is highly conserved across all species (Supplementary Figures 20 & 21) and the secondary structure distribution of the inner cavity largely remains constant (Supplementary Figure 22).

## 5 Discussion

Recent advances in biophysical techniques have led to remarkable insights into structural and dynamical properties of proteins, permitting an understanding of their morphology, nature as well as mechano-functional roles ^7^. Despite all these efforts, internal protein cavities, a predominant feature of most proteins, have been mainly ignored. This is in part due to their relatively inaccessible nature-making them experimentally challenging to characterize. Computational approaches can offer an ingenious solution to bridge this gap by speedily and accurately identifying residues lining the cavity, a prior step in their functional characterization. Several existing methods have attempted to access internal protein cavities (Figure 3). Yet, most of them are either severely limited in their usability (requiring user intervention such as MOLE *online*, Caver Web, HOLE, Molaxis) or functionalities (Caver Web). Given the variability in cavity shapes and the existing diversity of cavity detecting algorithms, it is likely that some methods may even perform better on specific orientations and conformation of cavities.

CICLOP, however, can characterize a versatile set of cavities (Supplementary Figure 6) like those contained within membrane-embedded channel proteins to cytosolic enzymes, chaperones, other cage like protein structures such as 3LOS and even multimeric viral capsids, which are massive in size (Figure 1h & i), with ease. For effective characterization of the cavity, CICLOP allows for both an automatic and manual mode of alignment of the input structure(s) with respect to the Z-axis which is essential in fine-tuning the detection process. Moreover, the atomistic resolution, an exclusive feature of our method (Supplementary Figure 7), allows the user to further calibrate the output results to serve their needs. Additionally, CICLOP can calculate the diameter as well as the volume of the pore along its length (Figure 1e & f), discern the overall nature of the cavity while also reflecting upon its secondary structure makeup (Figure 1d), compute its evolutionary conservation (Figure 1b & c) based on just the interface lining residues, aiding in a comprehensive functional annotation of the internal cavity.

We also performed extensive validation of our method and bench-tested it against existing methods in both qualitative as well as quantitative terms. The accuracy of CICLOP’s module to elucidate the diameter as a function of the cavity axis was validated by utilizing atomic coordinate distances in the crystal structure and was found to be within the mean error of 0.053 (Supplementary Figure 12). Furthermore, a comparison of the features of all the notable existing methods and CICLOP was tabulated (Figure 2). It is evident that CICLOP not only encompasses all the pre-existing methods but also provides some novel analytical implementations. Next, to understand the accuracy and precision of all the methods, we performed a quantitative in-silico experiment on cavities of three diverse proteins, a cytosolic kinase involved in the regulation of circadian rhythm (1TF7), a cytosolic chaperonin complex involved in protein folding (1AON) and a ligand-gated ion channel found at membrane interface (6V0B). This experiment highlighted the superior performance of CICLOP, where it exceeded other existing tools in terms of accuracy and precision of detection (Supplementary Figure 9, 10 & 11). Subsequently, a qualitative analysis on another heterogeneous set of proteins was performed against the two currently leading methods, Porewalker and MOLE *online*. This evidently showed that CICLOP could characterize a vivid set of cavity morphologies efficiently, while detection by both PoreWalker and MOLE*online* were misplaced such that they detected several residues visibly on the outer surface (Figure 3 and Supplementary Figure 14).

As a case study to exemplify CICLOP’s utility, we investigated the ATP synthase protein’s structure. ATP synthase is an archetypical example of a protein cavity undergoing rapid structural transitions in order to generate ATP. The cavity enclosed within the F1 domain is asymmetric (formed by the alternative arrangement of an *α* and *β* subunit) and is yet accommodating of the irregular helical rotor (*γ* subunit) ^2^, even as it moves ^9^. According to the rotary catalytic mechanism of ATP generation, the movement of *γ* subunit at every 120° steps is resultant in structural changes which are manifested in the catalytic sites (formed by the interface of *α* and *β* subunit) ^5;2^. This would only be possible through a physical interaction between residues of the rotor and the cavity interface. The intricate diameter profile of the sub-states generated by CICLOP not only illustrates these conformational changes (which are also magnified in the volume profile) but also showcases instances during the state transition where *γ* subunit is in very close proximity with the cavity interface (Figure 4e, Supplementary Figure 16 & 23). This intimacy between the two subunits is indeed expected, and such an interaction can warrant two physically disjointed units of a protein to work in sync ^17^. In the case of ATP synthase, the orientation of the bifurcated and slightly bent *γ* subunit towards a cavity interface might push the adjacent residues outwards to make space to accommodate it. This movement could in-turn lead to either the generation of a nucleotide-binding site on the outer surface such that its catalytic residues are now poised to act or abolish an existing site such that the bound nucleotide may be released. This is also depicted in Supplementary Figure 17b & c, where conformational changes have led to the appearance of a newer cluster of residues with one or more of their atoms facing the cavity during sub-state transitions. Elucidation of such cyclic regulatory mechanisms involving an otherwise inaccessible region such as a cavity is only possible through tools such as CICLOP that can detect even the microscopic changes occurring in protein cavity interface with extreme sensitivity and accuracy.

## 6 Conclusion

In summary, we have demonstrated our tool’s ability to quantitatively and qualitatively characterize the internal cavity of proteins. We have validated the results obtained by our tool using several in-silico methods. We have benchmarked our tool against existing methods in the field to find that the analysis provided by CICLOP are more sensitive, precise, and accurate. We have also demonstrated the use of various modules offered by our tool and how they can facilitate inference of biological functions. In the future, we plan to implement this algorithm on dynamic structures, such that it can identify and characterize changes in cavities not just spatially but temporally as well. We believe that the atomistic detail provided by CICLOP will have a significant application in the field of structural biology in evaluating protein structures, identifying solvent-accessible surfaces, protein oligomerization studies, understanding chaperone-assisted protein folding as well as functional characterization of channels. The method is available at https://ciclop.raylab.iiitd.edu.in.

## Acknowledgements

The authors would also like to thank the HPC facility of IIIT Delhi for the computational facility.

