## Supplementary file for "CICLOP: A Robust, Faster, and Accurate Computational Framework for Protein Inner Cavity Detection"

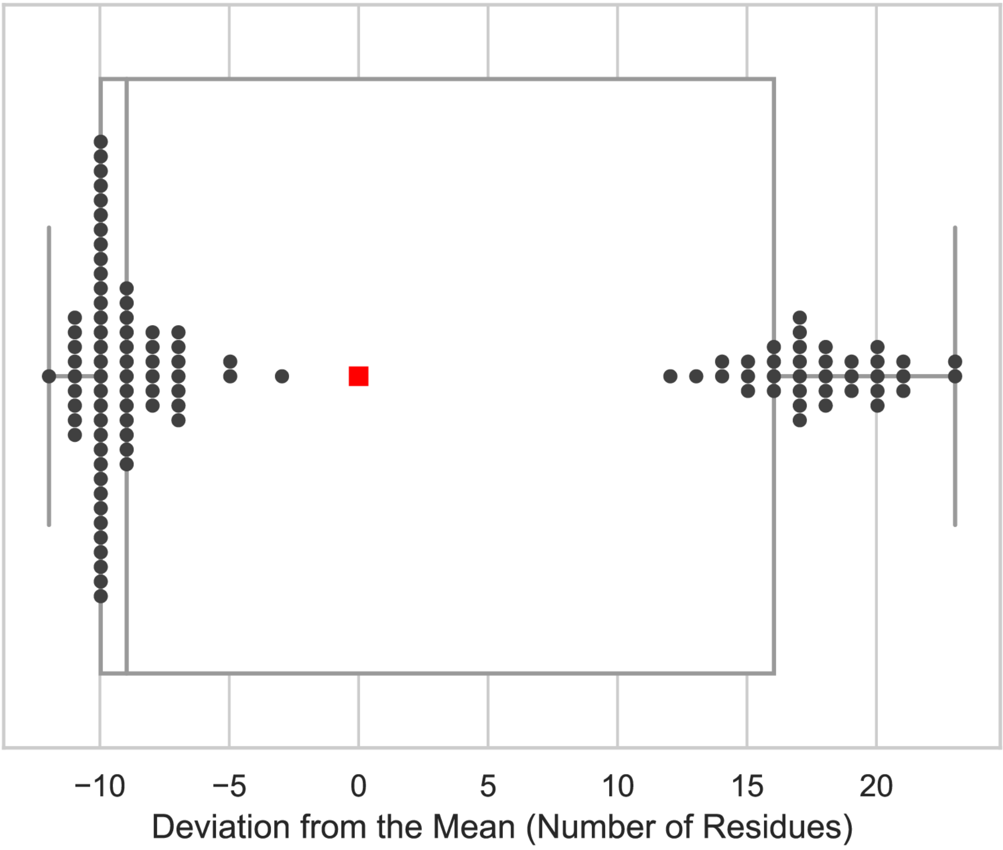


Figure 1: **Effect of the degree of rotation of the submitted structure imparted on CICLOP's performance in detecting residues on the inner surface.** The box plot shows the deviations observed from the mean number of residues detected on rotating protein structure(PDBID:1AON) every 10°. The observed deviations in rotational variance across the three planes lie corresponds to be within ±0.05% with respect to the total inner lining residues.

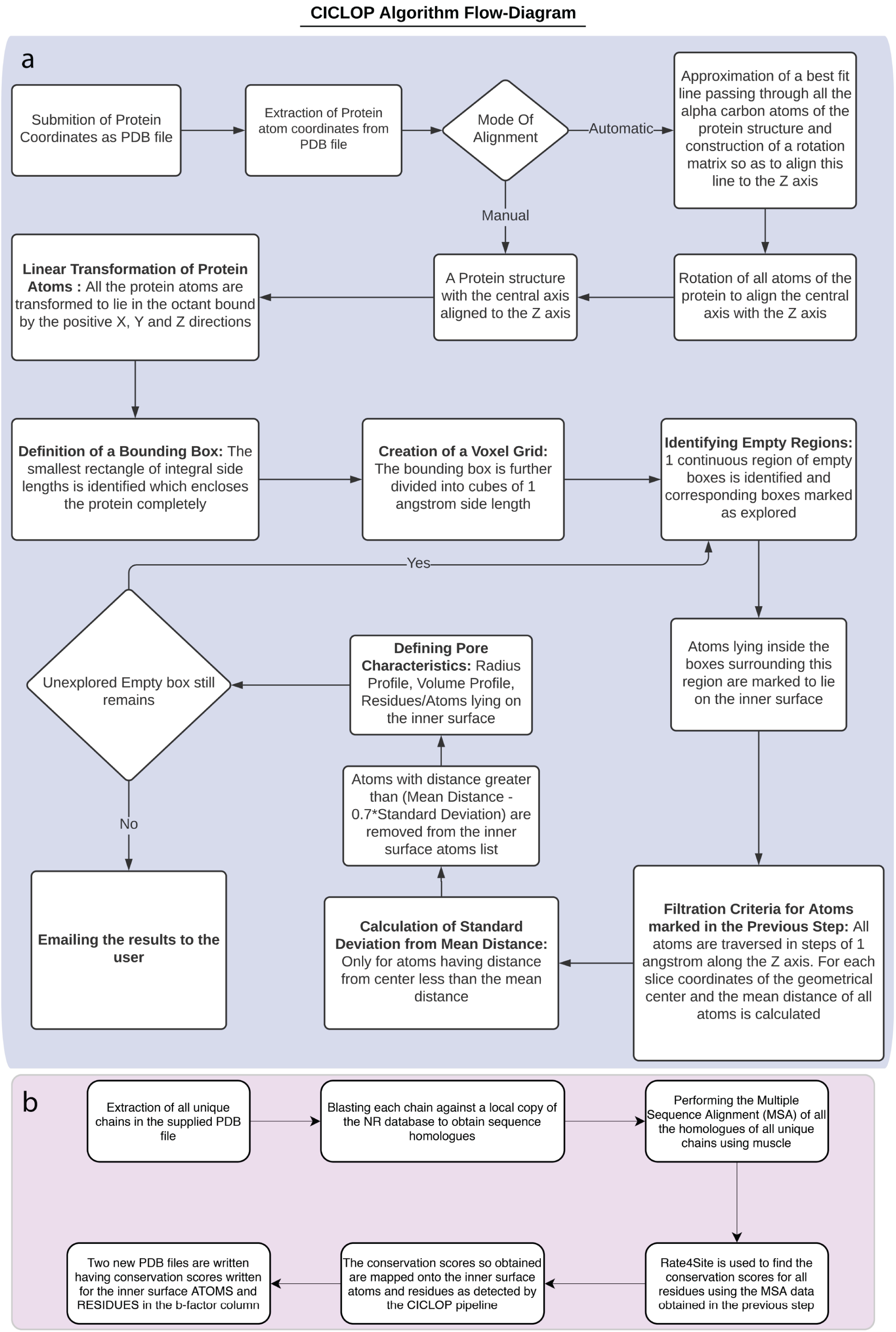


Figure 2: **CICLOP Workflow Diagram: a.** The basic process followed by CICLOP to find the inner cavity of a crystal structure in order to identify and characterize the residues lining the detected cavity. The volume and diameter profiles generated are also displayed. **b.** The methodology followed by CICLOP to calculate the conservation scores for the crystal structure as well as the cavity lining residues.

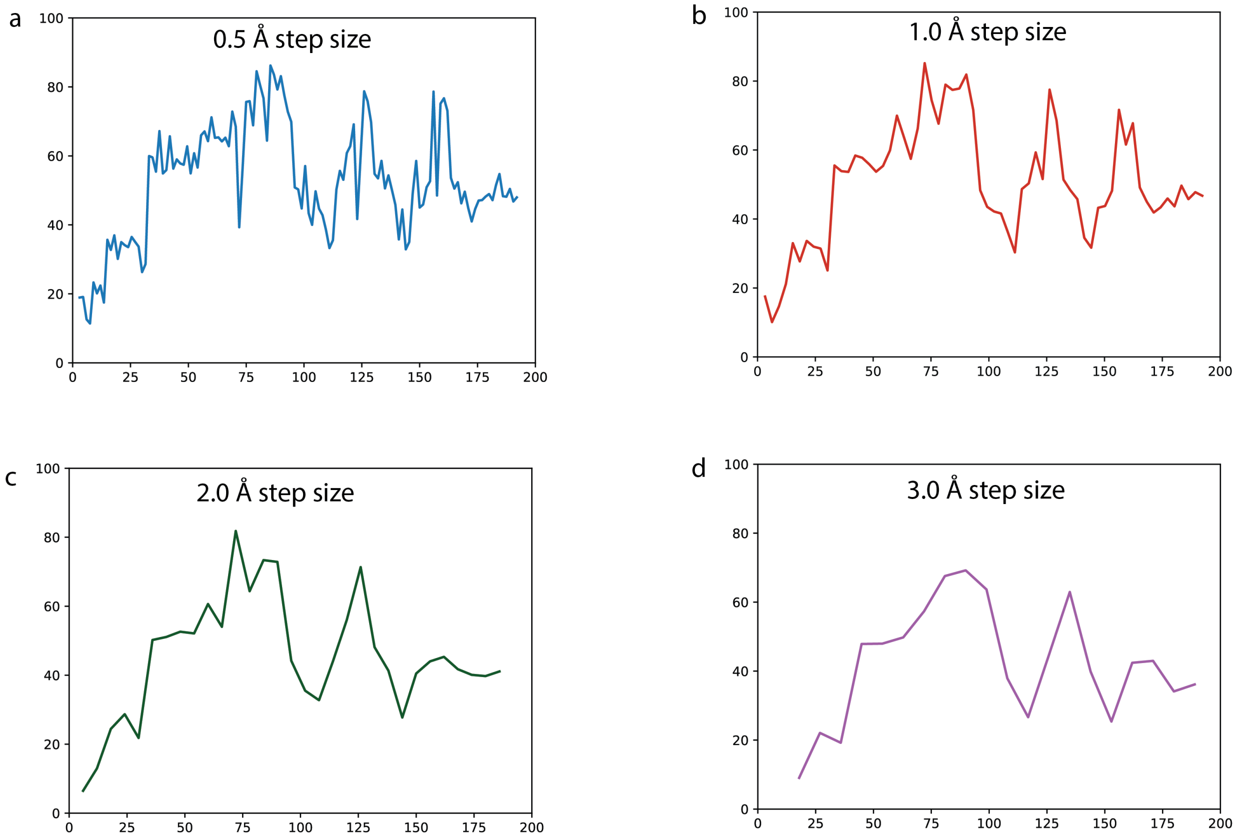


Figure 3: **Comparison of the diameter profile generated by CICLOP at different step size**

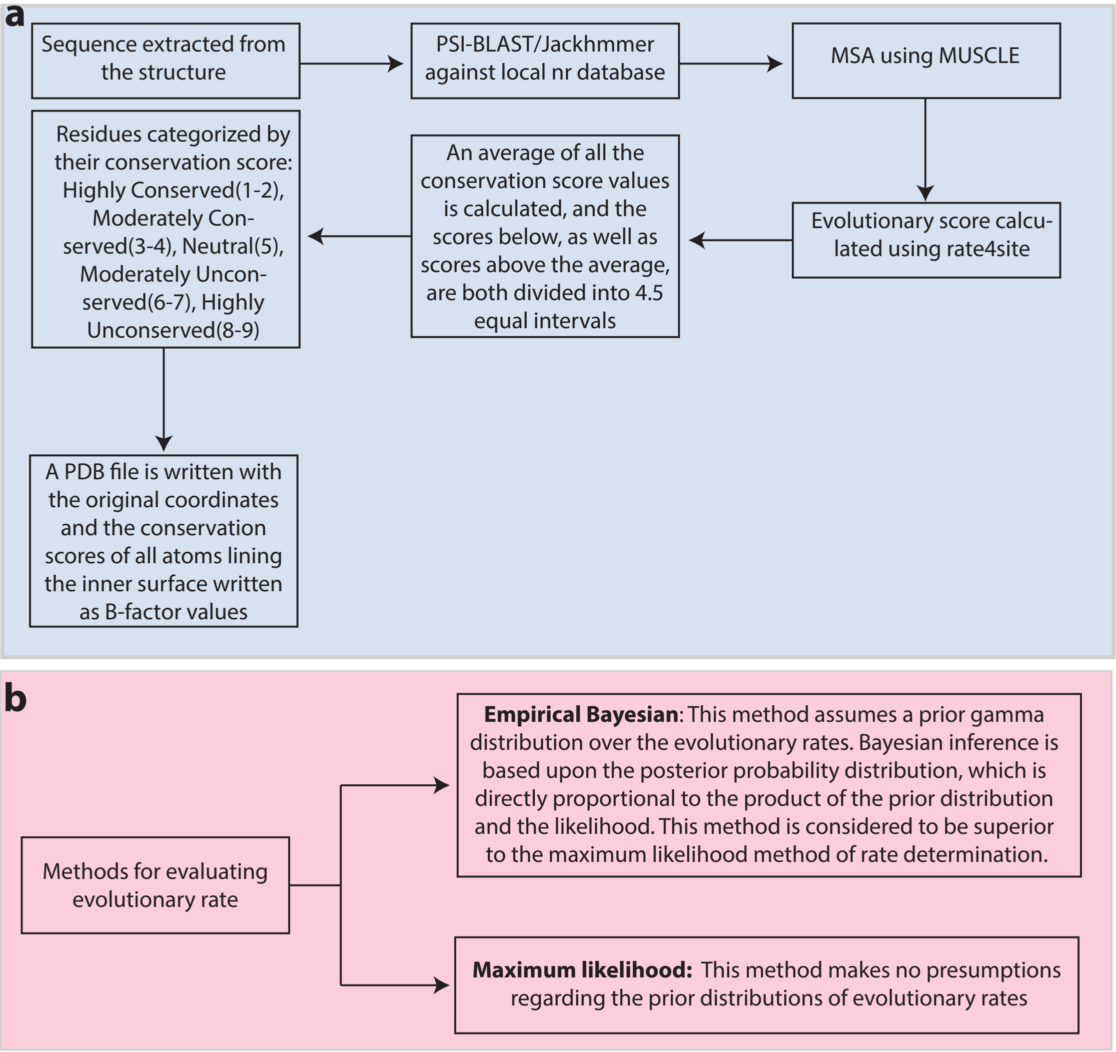


c

| Evolutionary Model | Description |
|---|---|
| JC amino acids | This model assumes that amino acids occur in equal frequency and are substitutedin equal probability. This method is chosen by default |
| Dayhoff Model | The empirical model of amino acid substitution developed by Dayhoff, Schwartzand Orcutt (1978) assumes that all sites in a protein evolve independently. |
| JTT Model | A new and improved version of the Dayhoff model developed by Jones, Taylor and Thornton. This uses a much larger dataset to construct a replacement matrix as proposed in the Dayhoff model |
| REV Model | The is the general reversible Markov process model of amino acid substitution. For this model, the probability of the first amino acid changing to the second is kept the same as thep robability of the second amino acid changing to first |
| WAG Model | A comparatively new model, offering a better statistical fit to the data. This method may provide more accurate estimates of phylogenetic trees than other existing models |
| LG Model | This model is an improvement to the WAG model. It incorporates the variability of evolutionary rates across sites in the matrix estimation and uses a larger and more diverse database than BRKALN (which was used to estimate the WAG model) |
| cpREV Model | A model similar to the REV model but optimised for chloroplast proteins |

Figure 4: **Workflow for calculation of conservation scores using CICLOP:** a. Summary of the basic pipeline for calculating evolutionary score. All the unique chains from the input PDB file are extracted (in case the user does not submit a FASTA file for the same). A basic local alignment search is performed on each unique chain in the PDB file against a local copy of the nr-database of proteins. Other protein sequences similar to the reference sequence are obtained using the PSI-BLAST package from the NCBI BLAST suite. A multiple sequence alignment using Muscle is performed amongst the reference sequence and the other similar sequences obtained from PSI-BLAST. Alternatively, it is also possible to query the target sequence against the Swissprot database using jackhmmer to obtain sequences distantly similar to the target. The rate4site method is applied to the resulting MSA file to calculate the evolutionary scores. b. Methods for estimating evolutionary rate. c. The seven substitution models offered by CICLOP used to infer the evolutionary conservation scores.

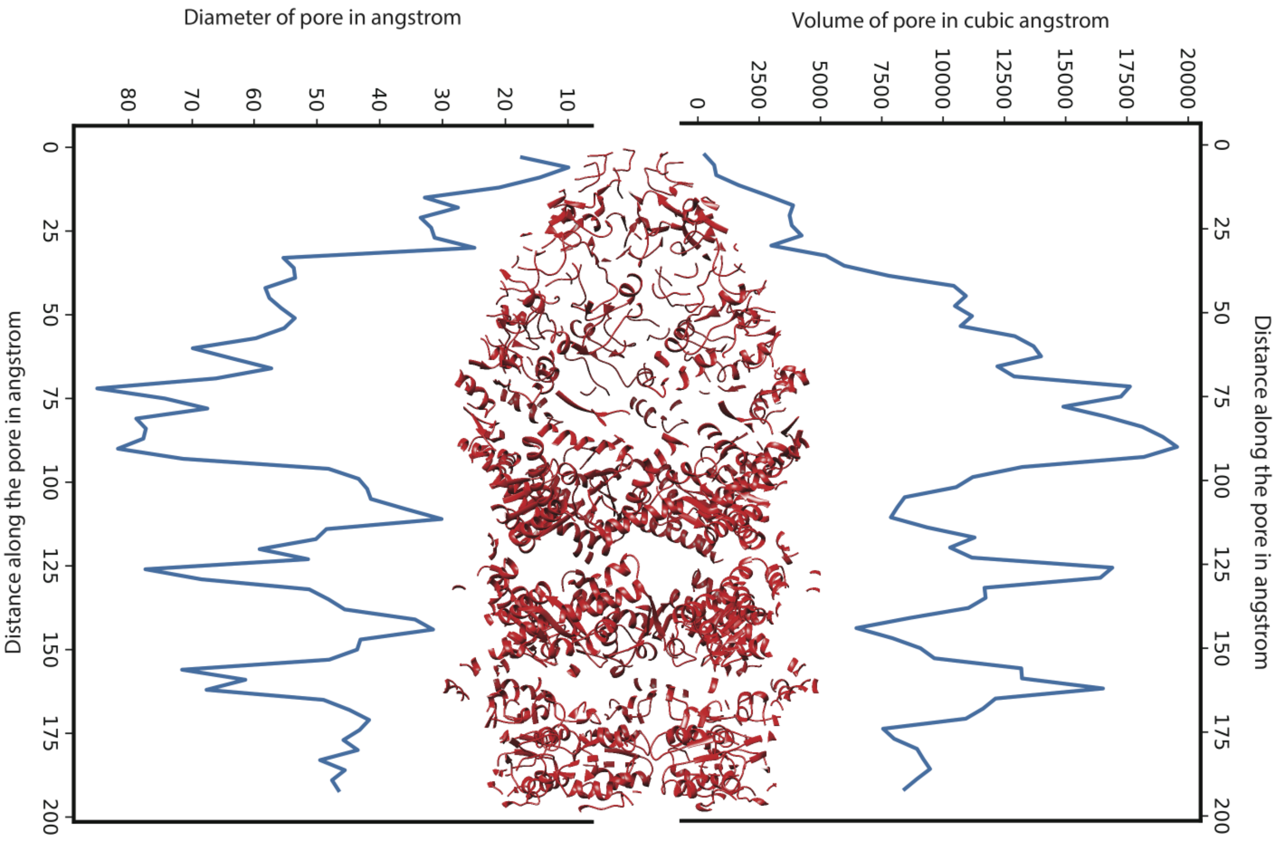


Figure 5: **Diameter and volume profiles produced by CICLOP as a function of pore length:** Diameter Profile (Left) and volume profile (Right) created by CICLOP along with the central cavity of the asymmetric GroEL/GroES/ADP complex (PDBID: 1AON).

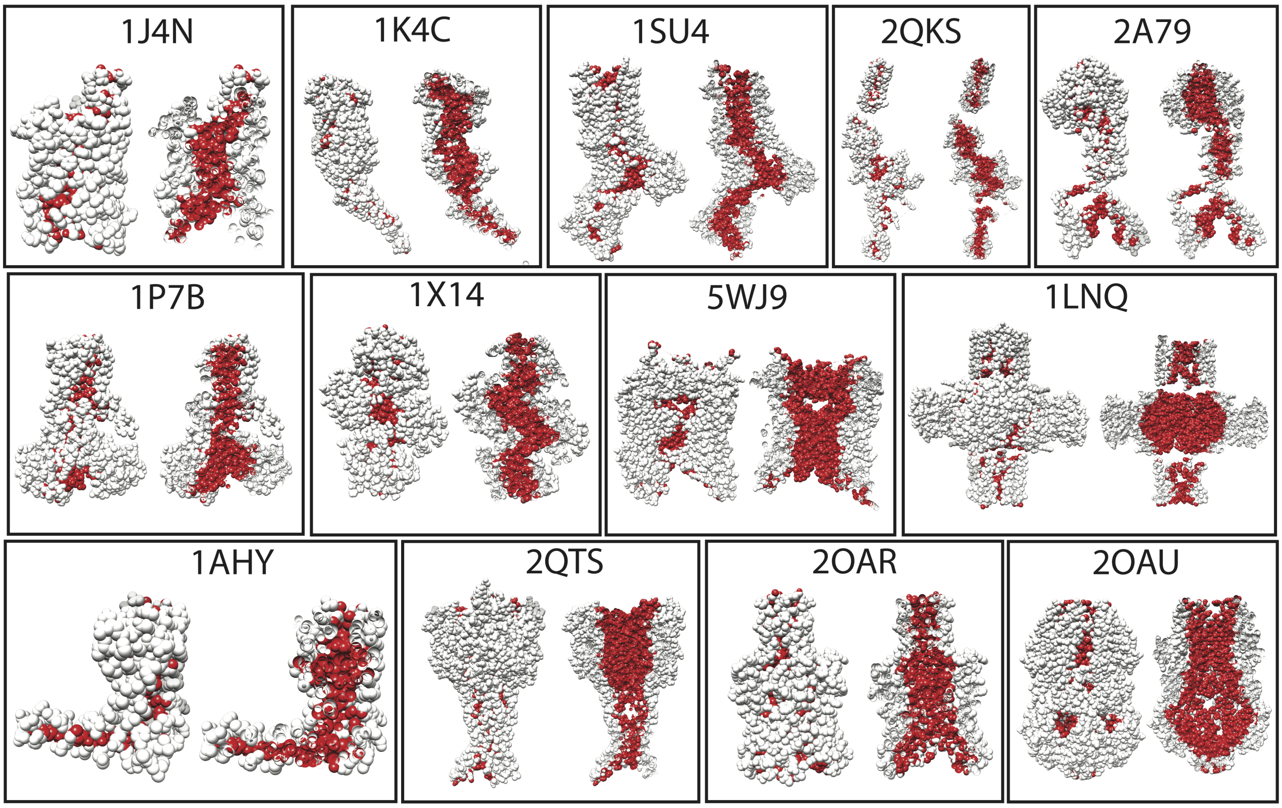


Figure 6: **Characterization of the inner cavity of proteins varying in size and structure using CICLOP.** Proteins ranging in cavity morphology, function as well as sub-cellular localization(cytosolic as well as transmembrane) were chosen to demonstrate CICLOP's usefulness in characterizing internal cavities. The complete surface (gray) as seen from the outside of the protein (left) as well as a longitudinal section of the protein (right) with the internal cavity identified by CICLOP (red). 1J4N, 1K4C, 1SU4, 2QKS, 2A79, 1P7B, 5WJ9, 1LNQ, 2QTS, 2OAR, 2OAU are transmembrane channel proteins while 1X14 and 1AHY are cytosolic enzymes.

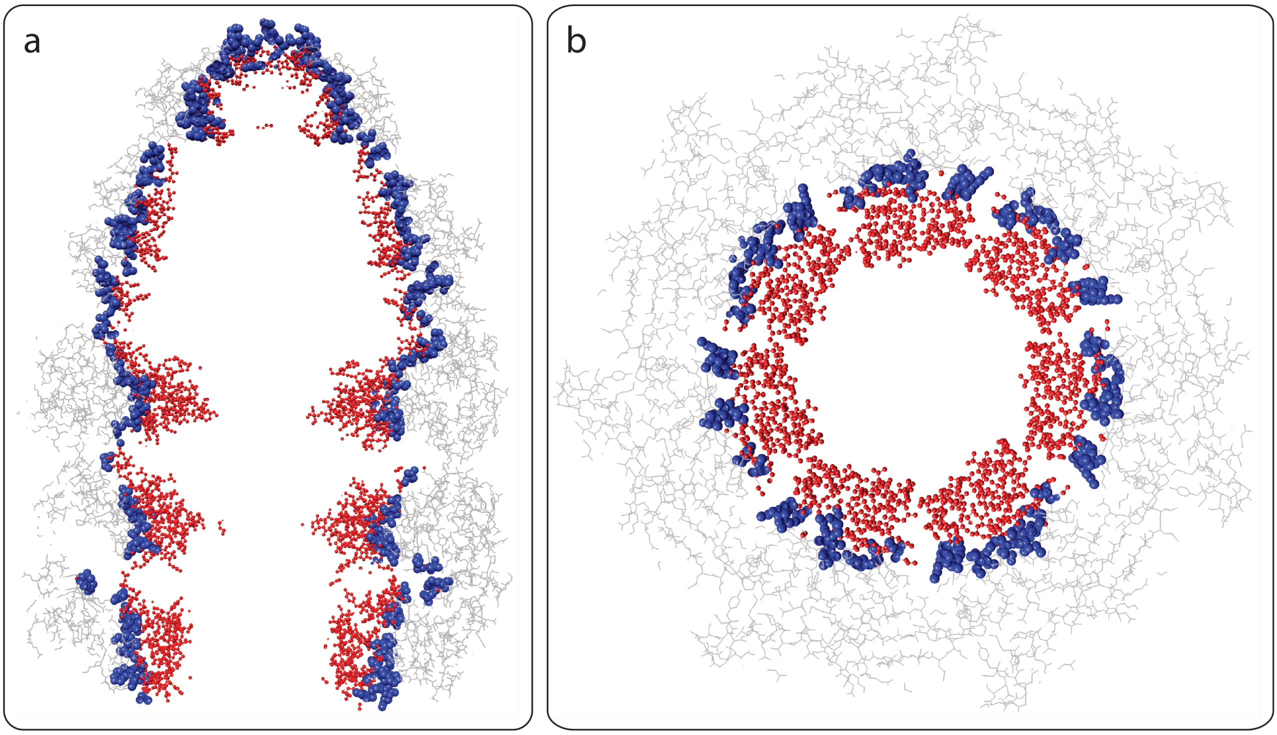


Figure 7: **Atomistic resolution of cavity detection. a.** Longitudinal section and **b.** Transverse section of the asymmetric GroEL/GroES/ADP complex (PDBID: 1AON). The bulk protein residues(gray wires) with residues that have 25-75% of their atoms detected to lie on the inside surface (blue). Residues with 100% atoms lining the cavity are marked in red.

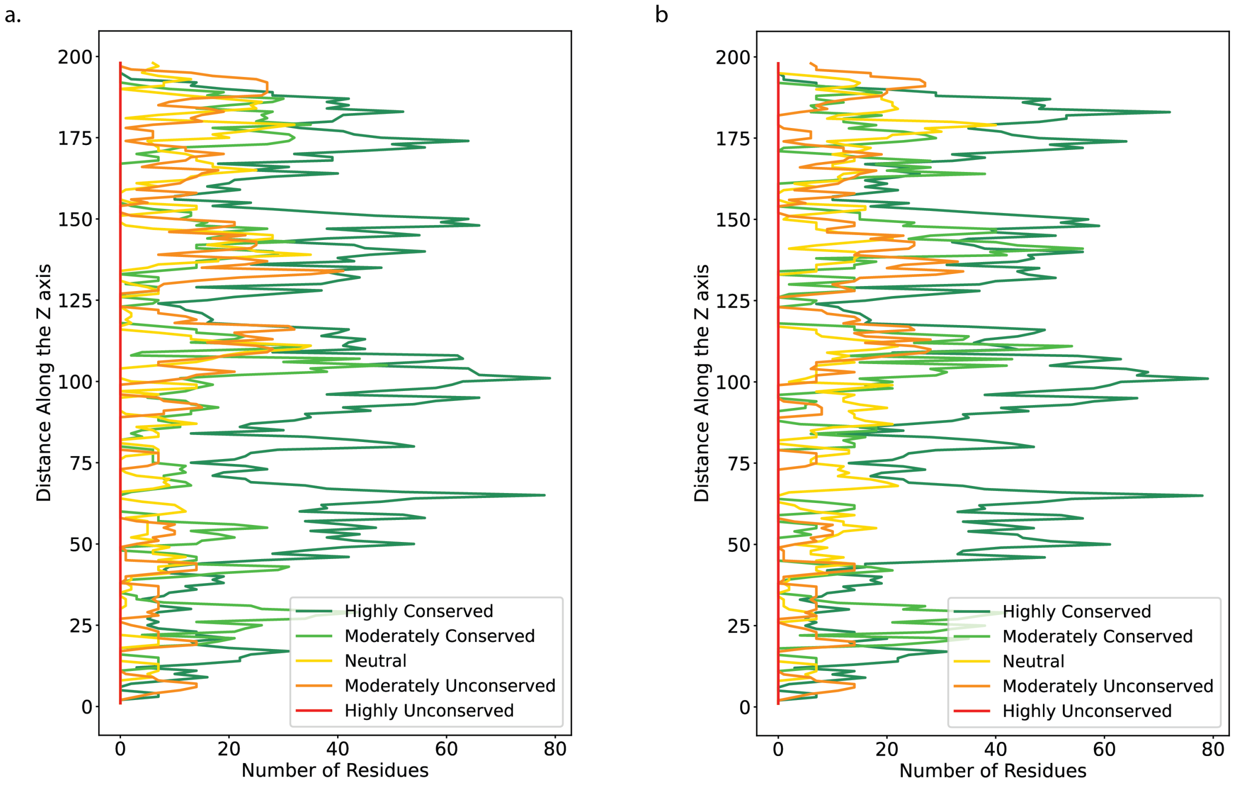


Figure 8: **Conservation profile of PDB ID:1AON generated by a. Psi-blast b. jackhmmer**

a

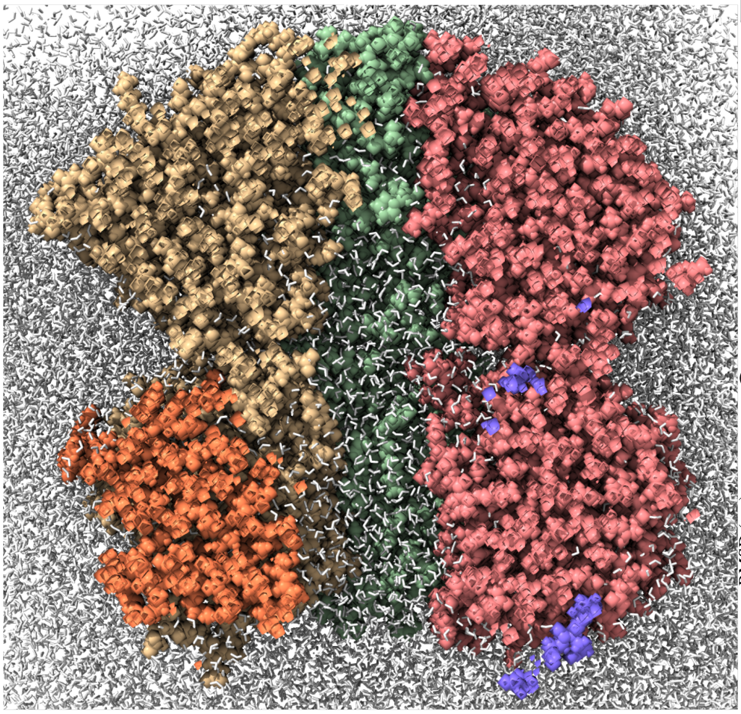

b

| Simulation Details @298K | | | |
|---|---|---|---|
| # of protein atoms | # of water atoms | # of ions | Total atoms (system size) |
| 45,798 | 244,227 | 30 | 2,90,055 |

c

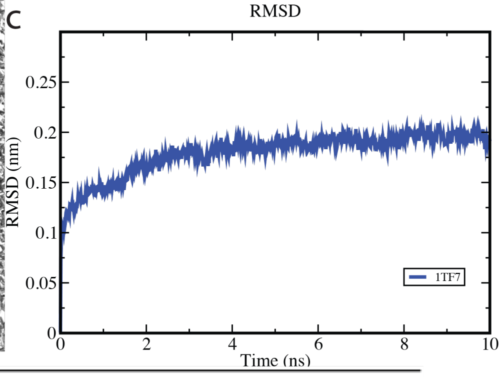


d

| | True Positive | False Positive | True Negative | False Negative | Accuracy | Precision |
|---|---|---|---|---|---|---|
| CICLOP | 85.58% | 5.01% | 94.98% | 14.41% | 91.52% | 90.85% |
| PoreWalker | 17.88% | 3.97% | 97.16% | 82.11% | 68% | 72.3% |
| MoleOnline | 31.64% | 11.38% | 88.67% | 68.35% | 67.69% | 61.79% |
| CaverWeb | 53.55% | 38.34% | 61.76% | 46.44% | 58.74% | 44.82% |

Figure 9: **Validation of the inner residues detected by CICLOP using MD simulation. a.** A snapshot of the longitudinal section of the simulated system. The protein is shown in surface representation (pastel colours) while the water molecules are shown in grey. **b.** The details of the simulated system for PDBID: 1TF7. **c.** The root mean square deviation (RMSD) plot of the protein across the simulation length. **d.** The table displays the accuracy and precision of CICLOP as well as other methods in detecting residues lining the inner cavity of 1TF7.

a

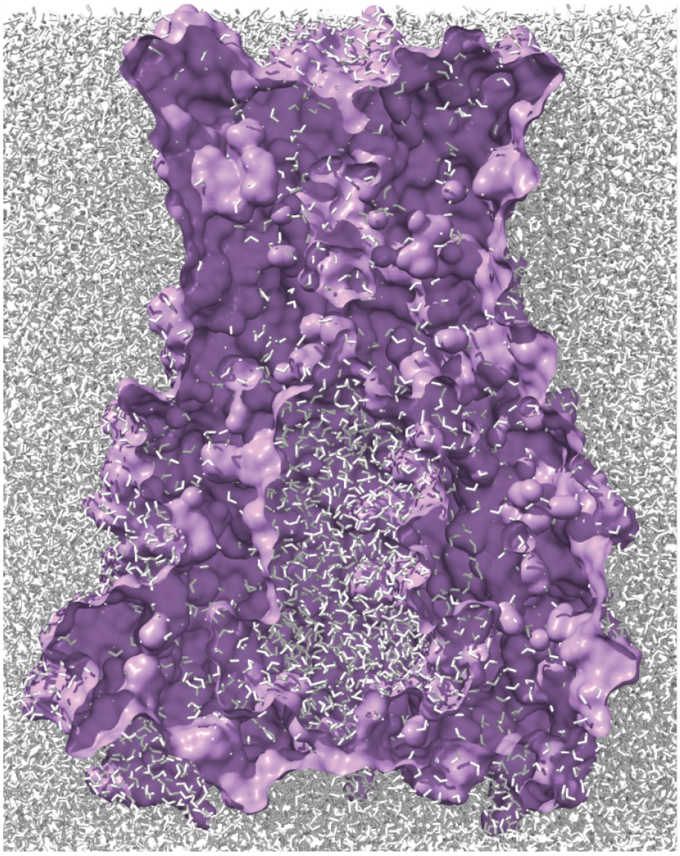

b

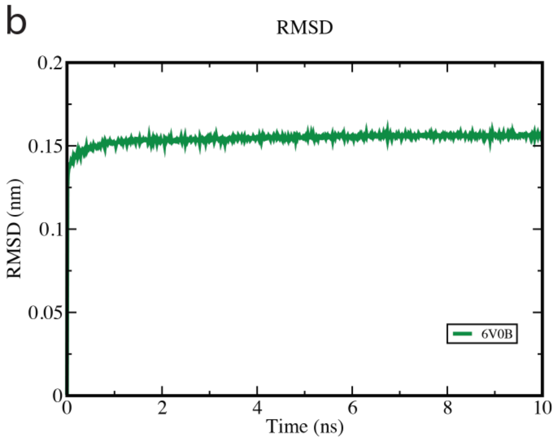


c

| Simulation Details @298K | | | |
|---|---|---|---|
| # of protein atoms | # of water atoms | # of ions | Total atoms (system size) |
| 24,835 | 229,335 | 30 | 254,200 |

d

| | True Positive | False Positive | True Negative | False Negative | Accuracy | Precision |
|---|---|---|---|---|---|---|
| CICLOP | 72.71% | 0.69% | 99.30% | 27.28% | 85.22% | 99.15% |
| PoreWalker | 29.62% | 0.55% | 99.44% | 70.37% | 62.48% | 98.36% |
| MoleOnline | 44.69% | 14.44% | 85.55% | 55.30% | 63.92% | 77.68% |
| CaverWeb | 9.87% | 4.58% | 95.41% | 90.12% | 50.13% | 70.79% |

Figure 10: **Validation of the inner residues detected by CICLOP using MD simulation. a.** A snapshot of the longitudinal section of the simulated system. The protein is shown in surface representation (purple) while the water molecules are shown in grey. **b.** The root mean square deviation (RMSD) plot of the protein across the simulation length. **c.** The details of the simulated system for PDBID: 6V0B. **d.** The table displays the accuracy and precision of CICLOP as well as other methods in detecting residues lining the inner cavity of 6V0B.

a

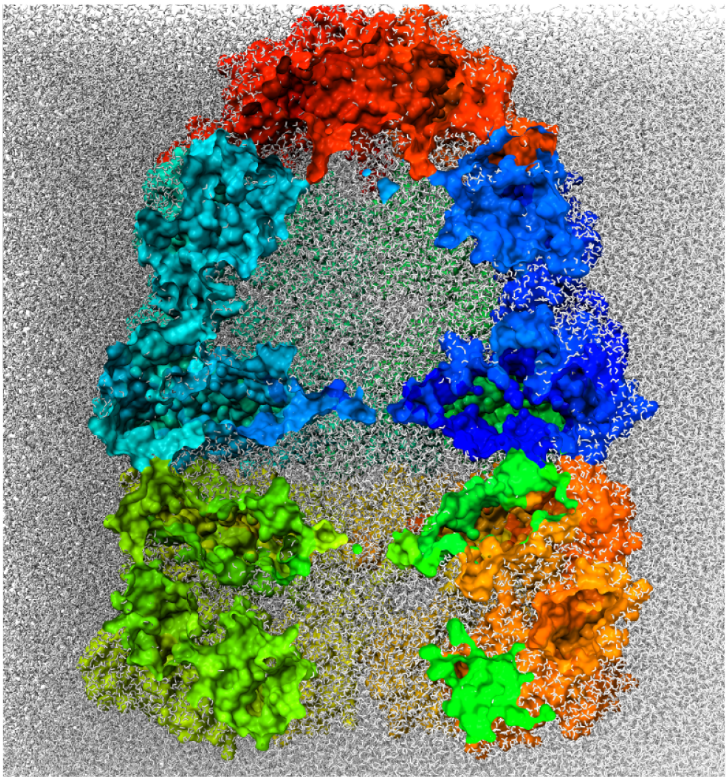

b

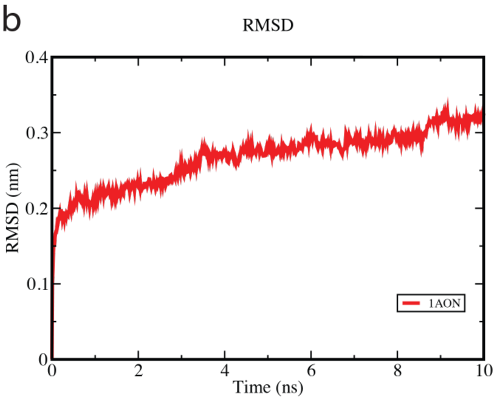


c

| Simulation Details @298K | | | |
|---|---|---|---|
| # of protein atoms | # of water atoms | # of ions | Total atoms (system size) |
| 120,120 | 1,064,094 | 273 | 1,184,487 |

d

| | True Positive | False Positive | True Negative | False Negative | Accuracy | Precision |
|---|---|---|---|---|---|---|
| CICLOP | 80.31% | 5.51% | 94.48% | 19.68% | 89.07% | 90.01% |
| PoreWalker | 29.44% | 5.08% | 95.15% | 70.55% | 70.04% | 78.16% |
| MoleOnline | 6.69% | 9.10% | 90.89% | 93.30% | 58.71% | 31.25% |
| CaverWeb | - | - | - | - | - | - |

Figure 11: **Validation of the inner residues detected by CICLOP using MD simulation. a.** A snapshot of the longitudinal section of the simulated system. The protein is shown in surface representation (rainbow) while the water molecules are shown in grey. **b.** The root mean square deviation (RMSD) plot of the protein across the simulation length. **c.** The details of the simulated system for PDBID: 1AON. **d.** The table displays the accuracy and precision of CICLOP as well as other methods in detecting residues lining the inner cavity of 1AON. Notably, Caver Web is unable to process this structure.

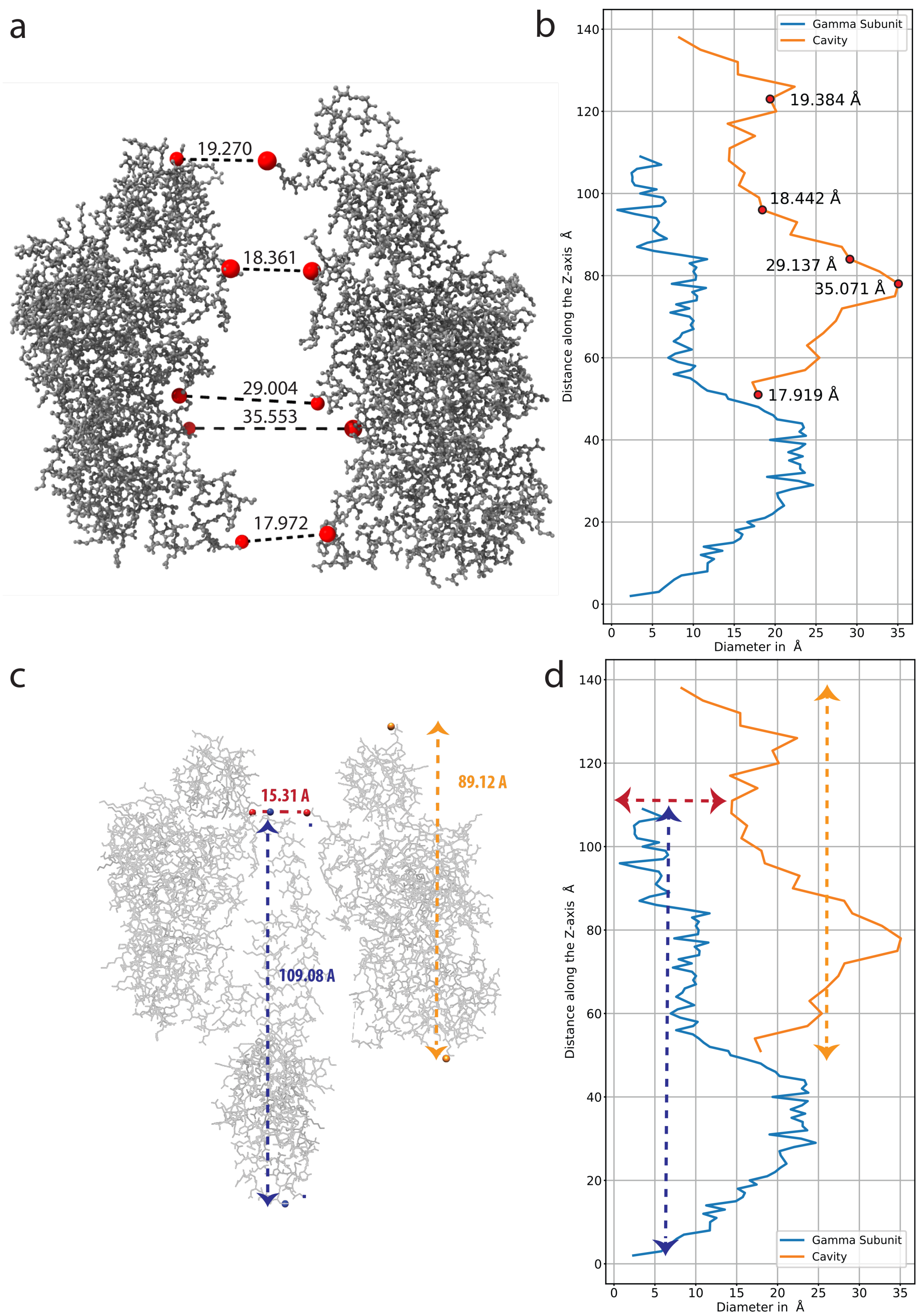


Figure 12: **Verification of the size of the cavity as indicated by CICLOP through manual annotation (using UCSF Chimera) a.** Validation of the diameter profile generated using CICLOP through manual annotation (using UCSF Chimera). Distances between atoms lying on the opposite ends of chains A and F of State 1A are marked (left) along with the corresponding points on the **b.** diameter profile generated by our tool. **c.** Validation of the size of the cavity. Distances between the atom with the minimum and maximum Z-axis value are marked (in yellow) for chain B (representative of the cavity) as well as chain G ( $\gamma$ subunit) of state 1A. **d.** The corresponding values are indicated in the diameter profile generated by CICLOP.

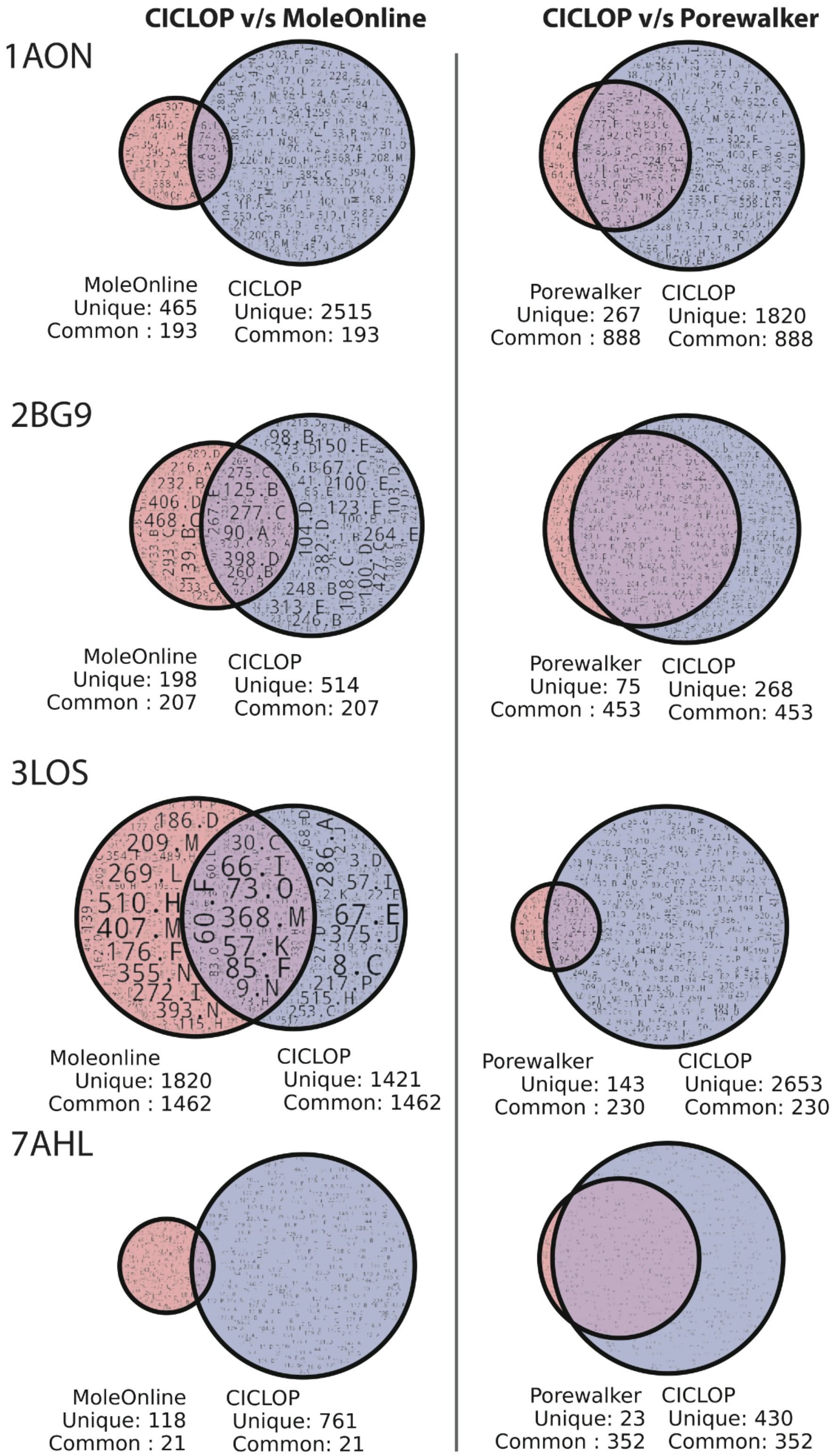


Figure 13: Quantification by venn diagram representation for the number of residues detected by CICLOP and other methods (PoreWalker and MOLE*online*) as shown in Figure 8.

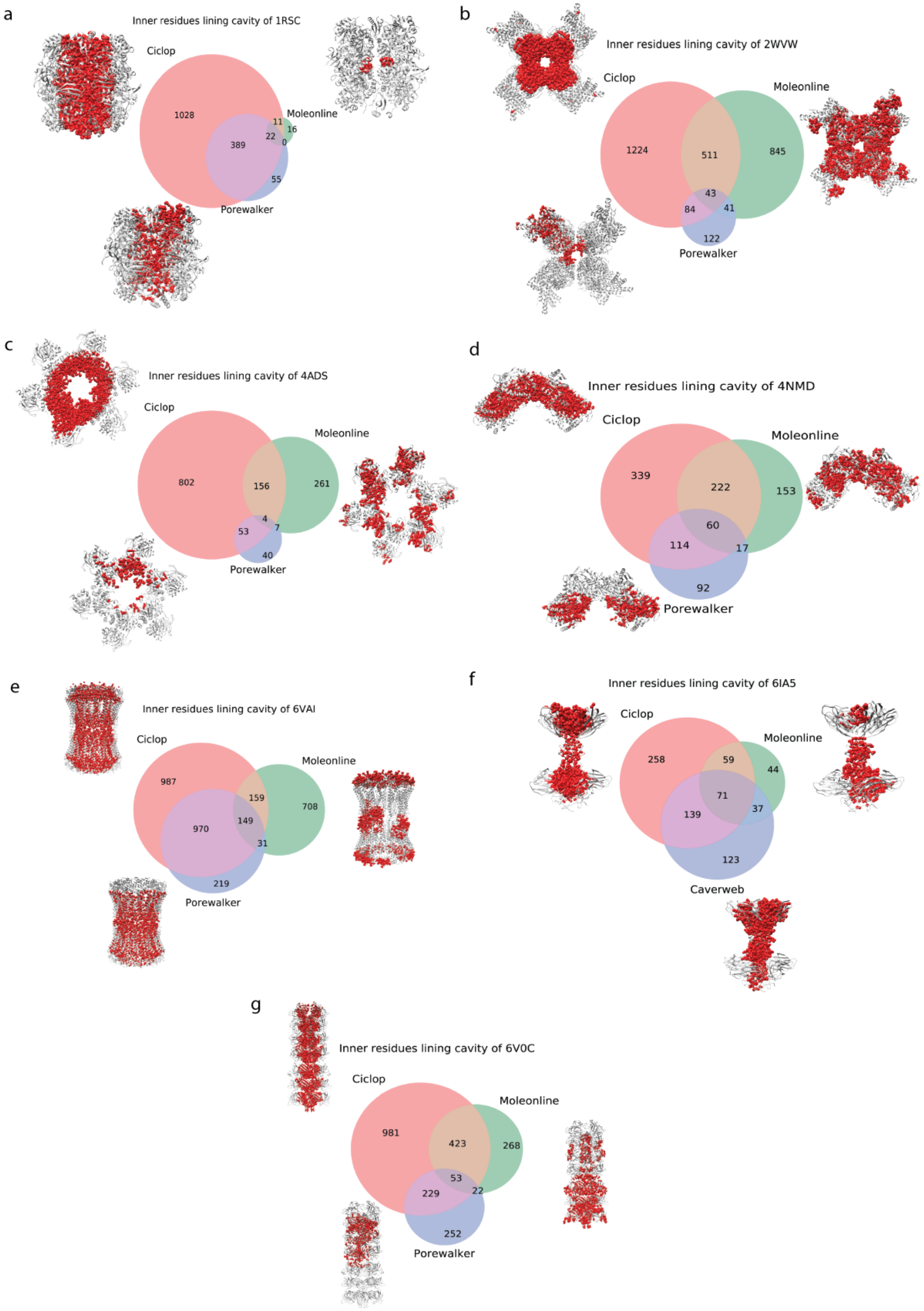
a
Inner residues lining cavity of 1RSC
Ciclop
Moleonline
Porewalker
1028
389
11
16
22
0
55
b
Inner residues lining cavity of 2WVW
Ciclop
Moleonline
Porewalker
1224
511
845
43
84
41
122
c
Inner residues lining cavity of 4ADS
Ciclop
Moleonline
Porewalker
802
156
261
4
53
7
40
d
Inner residues lining cavity of 4NMD
Ciclop
Moleonline
Porewalker
339
222
153
60
114
17
92
e
Inner residues lining cavity of 6VAI
Ciclop
Moleonline
Porewalker
987
159
708
149
970
31
219
f
Inner residues lining cavity of 6IA5
Ciclop
Moleonline
Caverweb
258
59
44
71
139
37
123
g
Inner residues lining cavity of 6V0C
Ciclop
Moleonline
Porewalker
981
423
268
53
229
22
252

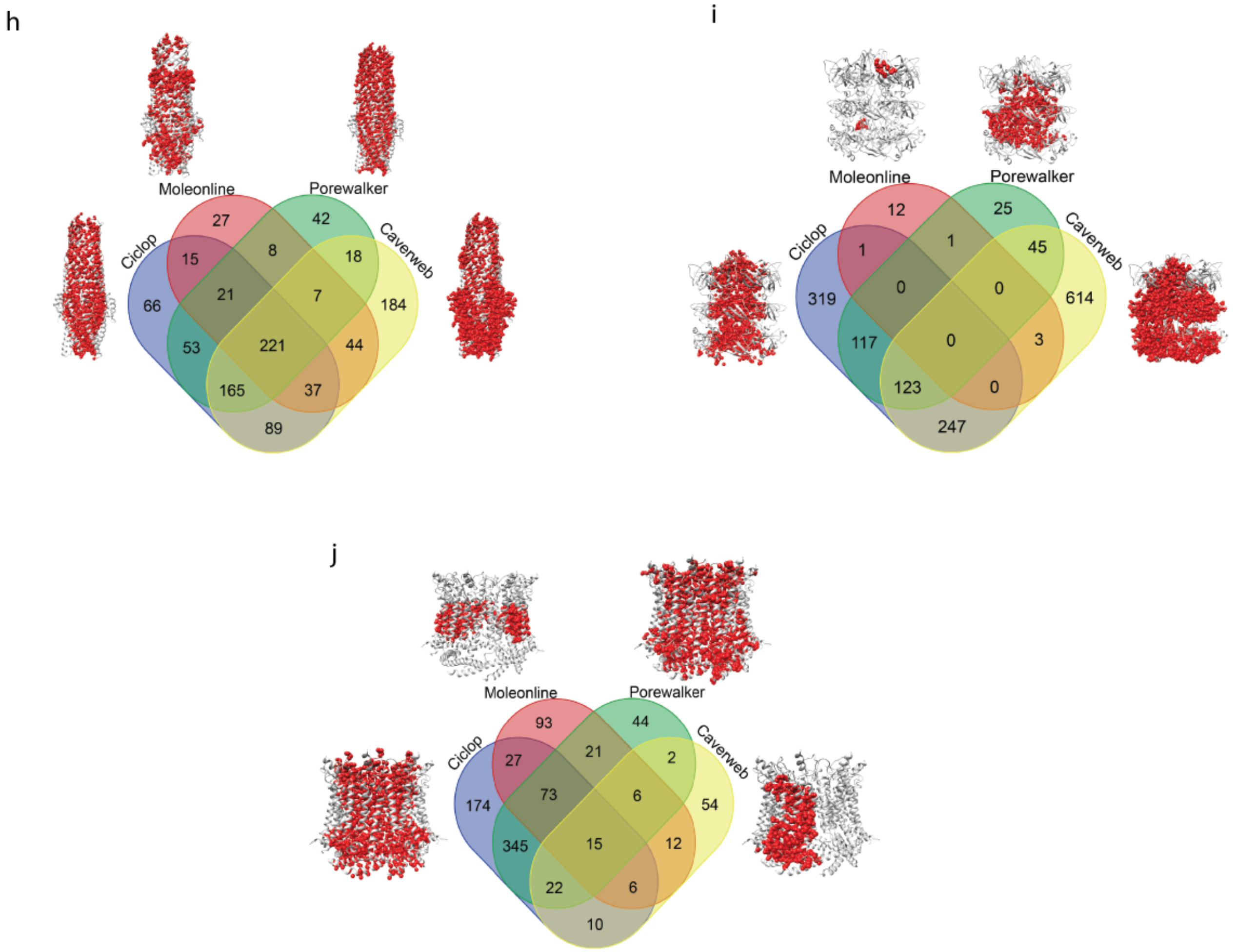


Figure 14: **Venn diagram showing the number of inner residues detected by tunnel detecting method on proteins of various size and cavity conformation.** The proteins are represented as gray ribbons and the residues detected on the inner surface are rendered as red spheres for easy visualization.

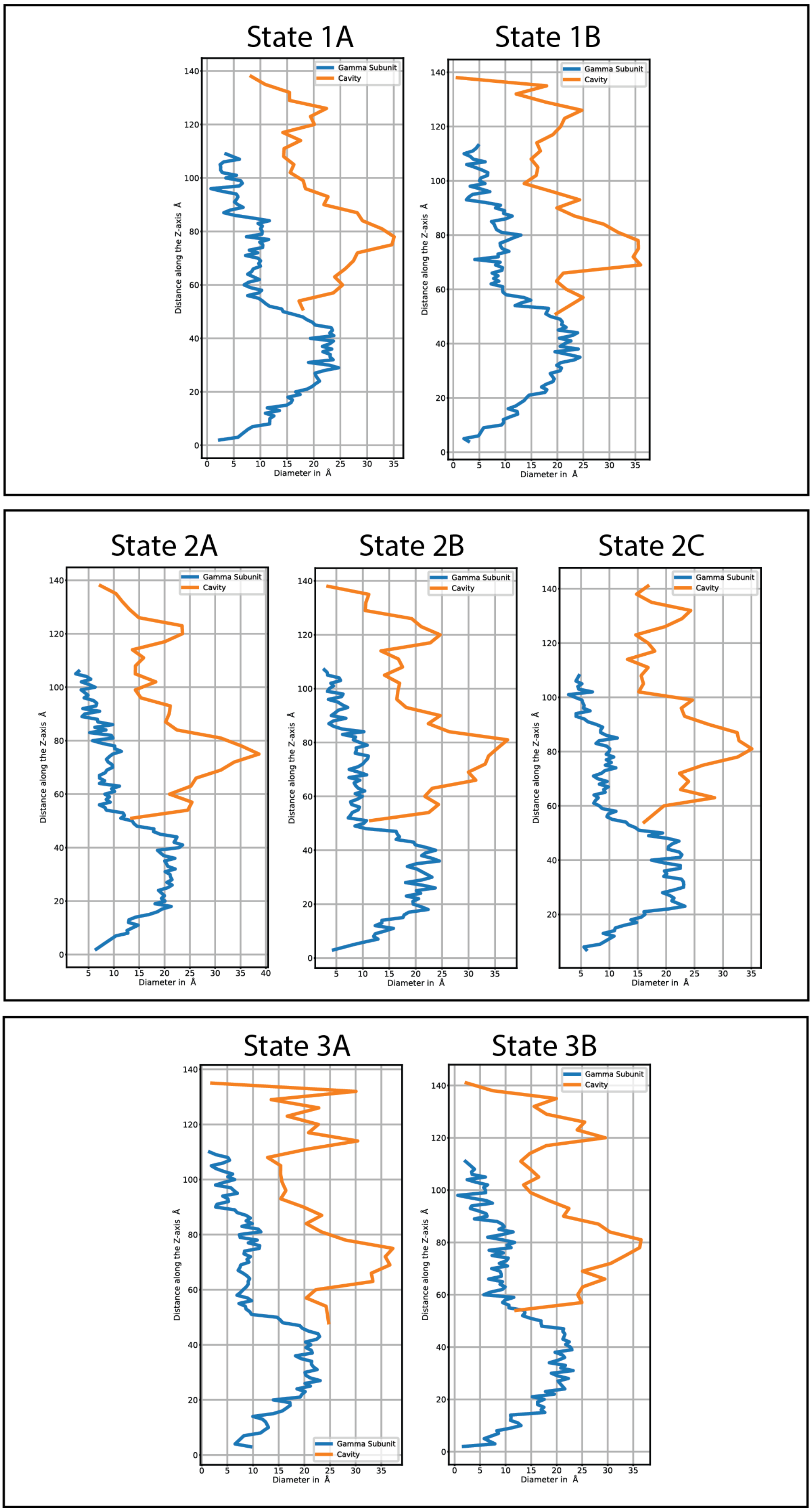


Figure 15: **Diameter profiles of the cavity of ATP synthase F1 domain during its catalytic rotation generated using CICLOP.** The figure highlights the heterogeneous changes in the size of the cavity due to conformational changes occurring during the rotation of $\gamma$ subunit. The diameter of the top and bottom part of the cavity fluctuates due to orientation of the rotor in the various substates.

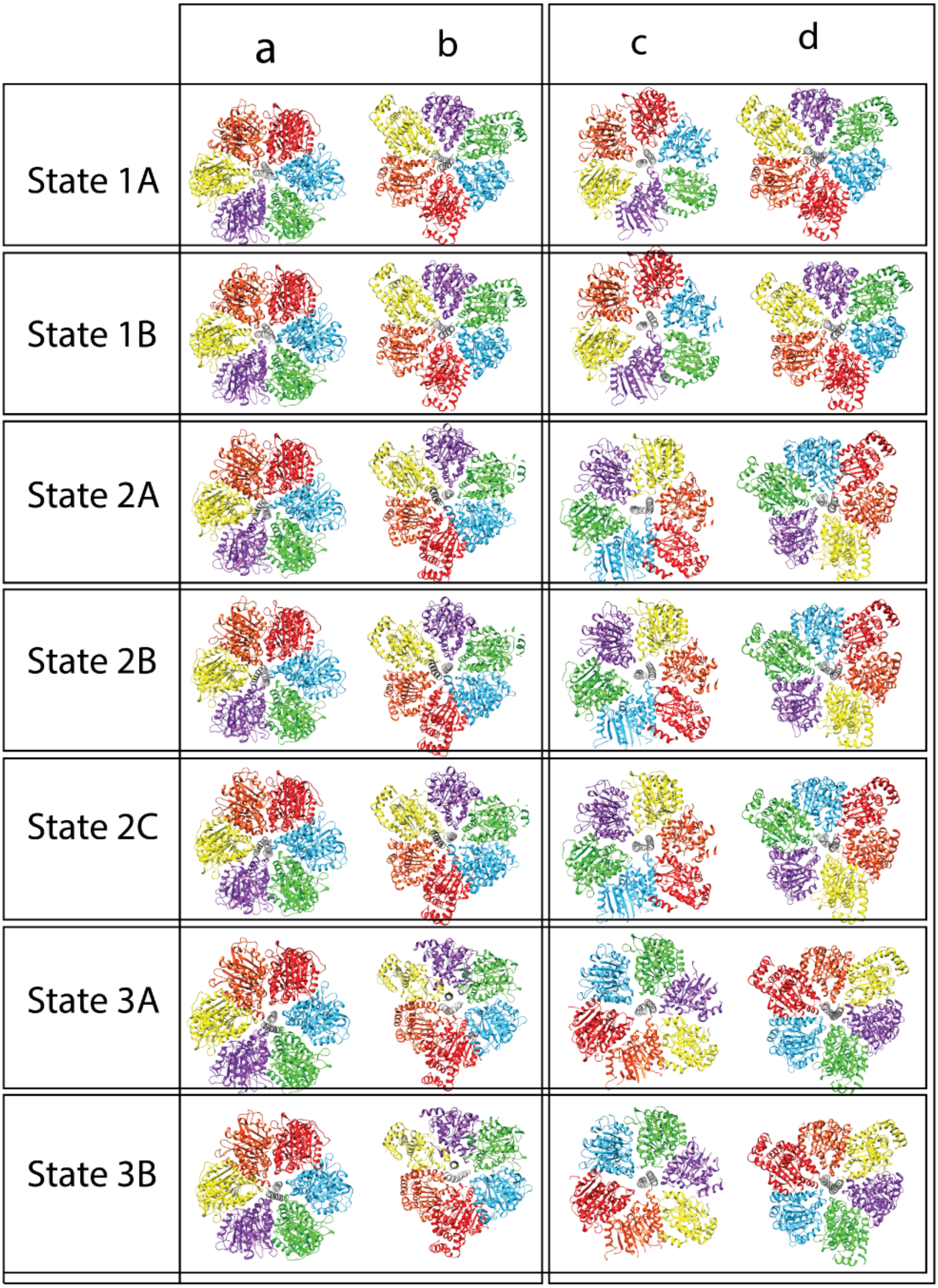


Figure 16: **Orientation of ATP synthase F1 domain during its rotary catalytic cycle.** Panel a and b depict the orientation of the $\gamma$ subunit or chain G (gray) with respect to the cavity. Panel c and d represent the orientation of the cavity with respect to chain G (where chain A, B, C, D, E, F are shown as red, yellow, green, blue, orange, purple respectively). Panels a and c are top views while b and d are bottom views of the cavity. The figure shows the orientation of $\gamma$ subunit towards the $\alpha$ $\beta$ interface. Each rotational state (1-3) is related to each other by the rotation of $\gamma$ by $120°$.

a

| State | Transitions | Residues present in Set B but not Set A (Set B - Set A) | Residues present in both Set B and Set A (A ∩ B) | Total residues detected by CICLOP | Percentage change in residues lining cavity during transitions |
|---|---|---|---|---|---|
| 1A | 3B to 1A | 1A - 3B = 282 | 647 | 929 | 30.35 % |
| 1B | 1A to 1B | 1B - 1A = 227 | 704 | 931 | 24.38 % |
| 2A | 1B to 2A | 2A - 1B = 143 | 808 | 951 | 15.03 % |
| 2B | 2A to 2B | 2B - 2A = 63 | 854 | 917 | 03.92 % |
| 2C | 2B to 2C | 2C - 2B = 77 | 861 | 938 | 08.20 % |
| 3A | 2C to 3A | 3A - 2C = 158 | 777 | 935 | 16.89 % |
| 3B | 3A to 3B | 3B - 3A = 96 | 843 | 939 | 10.22 % |
| **Residues common in all 7 substates = (1A ∩ 1B ∩ 2A ∩ 2B ∩ 2C ∩ 3A ∩ 3B)= 557** | | | | | |

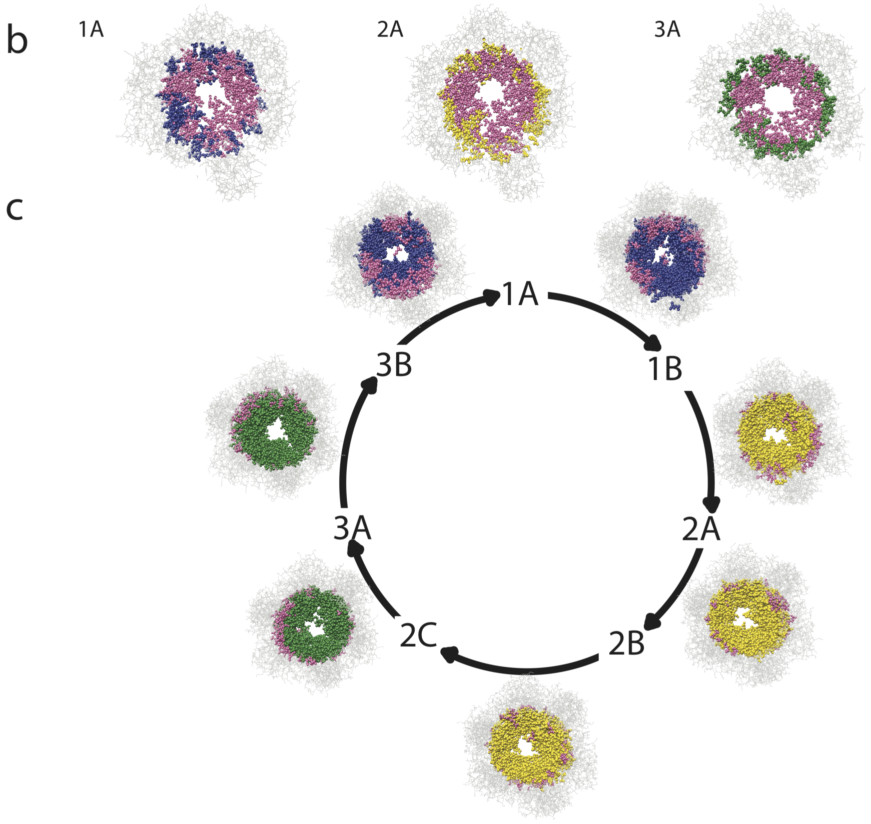


Figure 17: **Distribution of residues detected in the inner cavity of each sub-state of ATP synthase F1 domain occurring during ATP synthase catalytic cyclea.** Table displaying the quantified percent changes in residues lining the cavity of the F1 cavity during sub-state transitions. **b.** Top view of the cavity of each representative state during the ATP synthase rotary cycle. Residues facing the cavity as detected by CICLOP are coloured as blue, yellow and green for each state respectively while residues that are common in all the substates are coloured as pink.**c.** Top view of the cavity during each state transition of the catalytic rotation. Residues facing the cavity as detected by CICLOP are indicated in blue, yellow and green colours respectively for state 1, 2 and 3 respectively. Dynamic residues for each substate transition are marked in pink.

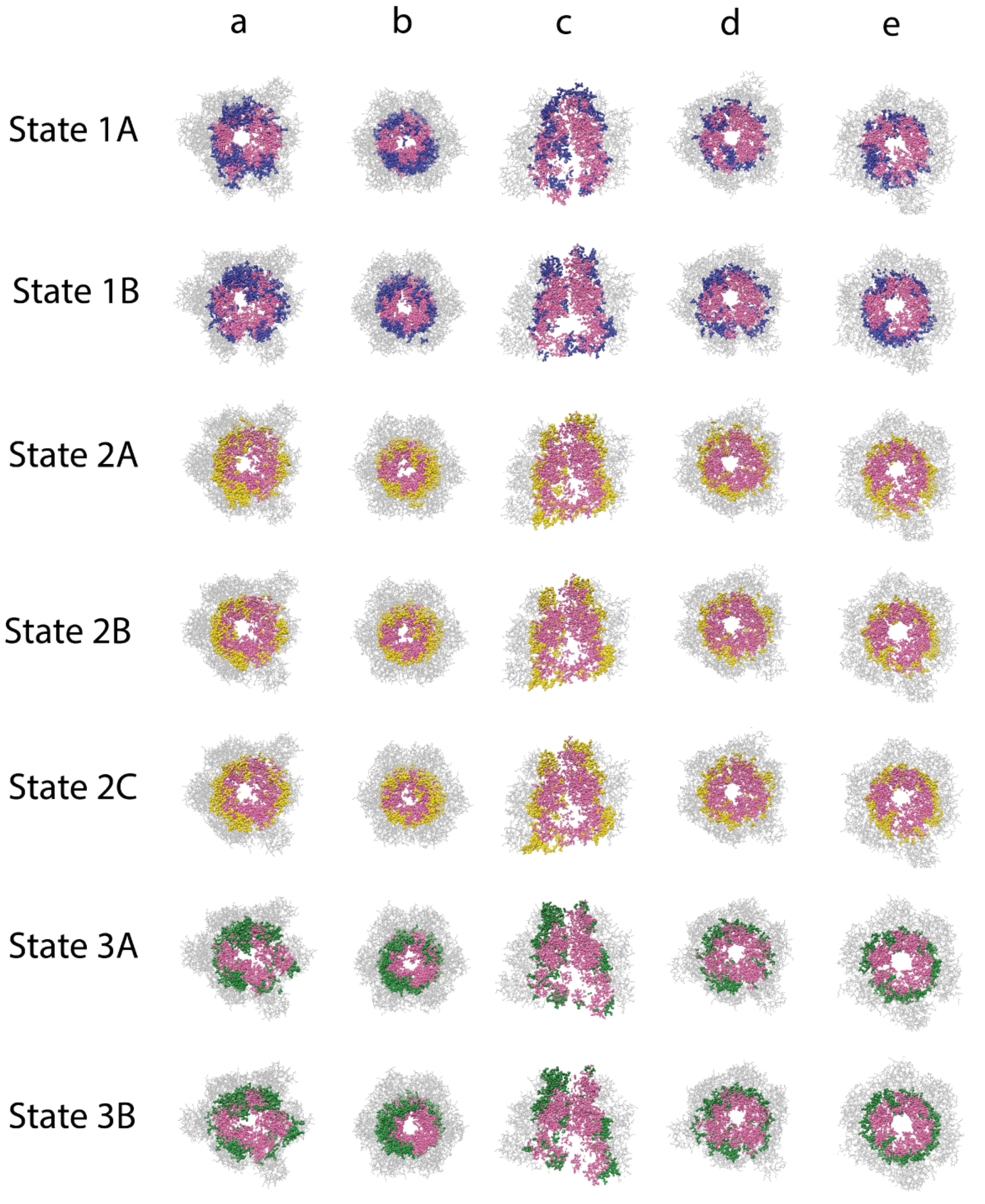


Figure 18: **Residues lining the inner cavity of F1 ATP synthase throughout its catalytic cycle.** The cavity is shown as gray wires while the residues detected on the inner surface are shown as colored spheres (blue, yellow and green for state 1 , 2 and 3 respectively). Residues that remain on the inside during the entire rotary cycle (present in all seven substates) are colored in pink. Panel a and b depict the bottom and top views of the cavity, c depicts a longitudinal section, while d and e depict a transverse section of the cavity as observed from the bottom and top respectively. The figure highlights that the residues immediately facing the cavity are well represented throughout all the substate transitions(pink), however during each substate transition, conformational changes lead to newer residues surfacing that now face the cavity (blue, yellow and green respectively for each state transition).

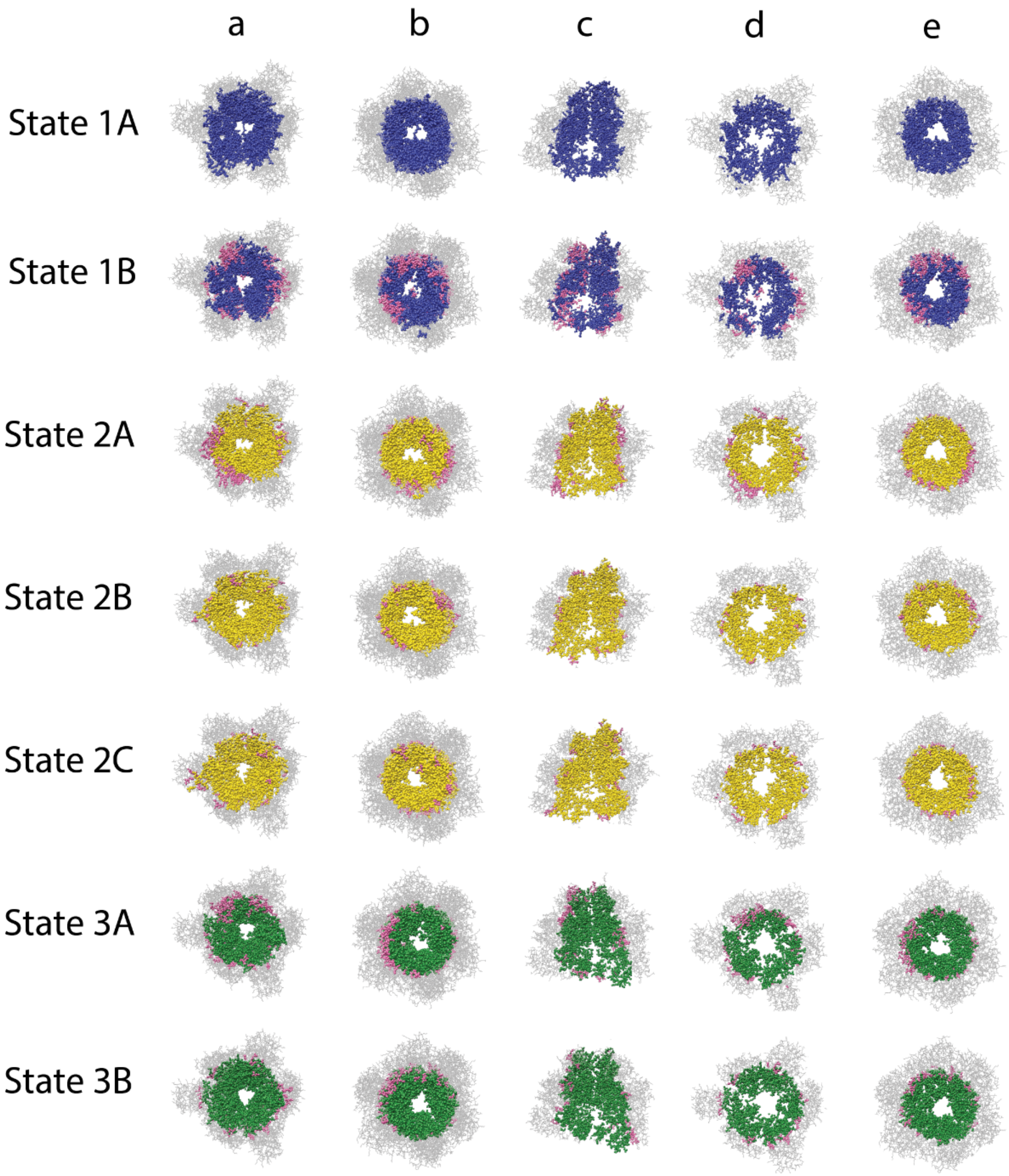


Figure 19: **Residues lining the cavity of F1 ATP synthase during each state transition.** The cavity is shown as gray wires while the residues detected on the inner surface are shown as colored spheres(blue, yellow, green for state 1, state 2, state 3 respectively). Newer residues that surface during a substate transition (new residues now facing the cavity) are shown in pink. Panel a and b depict the bottom and top views of the cavity, c depicts a longitudinal section, while d and e depict a transverse section of the cavity as observed from the bottom and top respectively. The figure highlights that the residues lining the internal face of the cavity (surrounding $\gamma$ subunit) always remain constant during state transitions (1-3) (blue, yellow and green), while residues that change lie diagonally opposite to each other(pink).

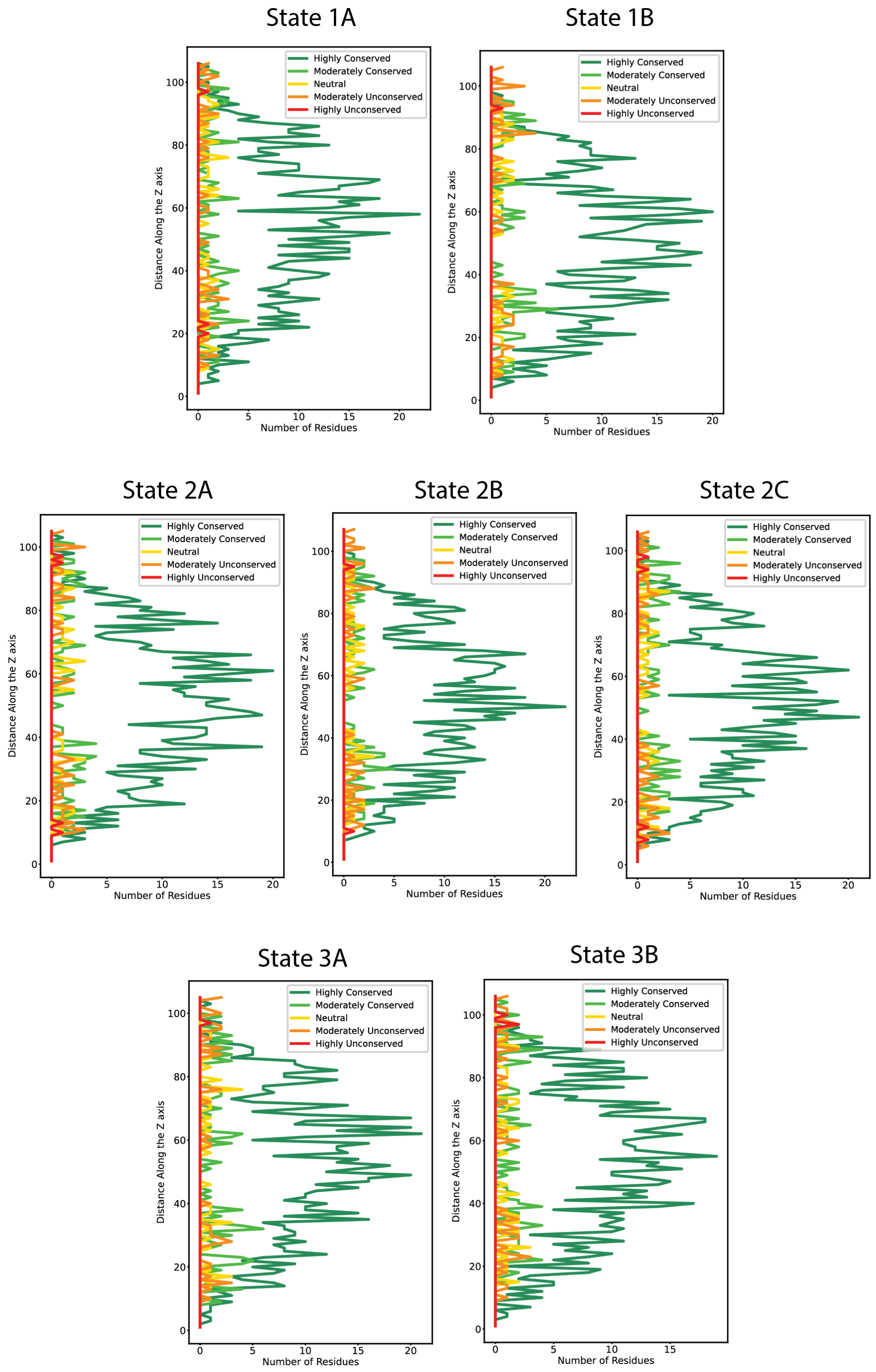


Figure 20: **Conservation profiles of residues lining the cavity of ATP synthase F1 domain generated by CICLOP in different states.** The residue-wise conservation frequency as a function of Z-axis length where the colours corresponding to CICLOP's scale is as follows: highly conserved = dark green, moderately conserved = light green, neutral = yellow, moderately unconserved = orange and highly unconserved = red. The internal cavity is lined by residues which are highly conserved across all species, while moderately-highly unconserved residues are scarcely present (mostly at the top of the structure). This figure reflects on the conserved function of F1 cavity of ATP synthases due to virtue of their structure.

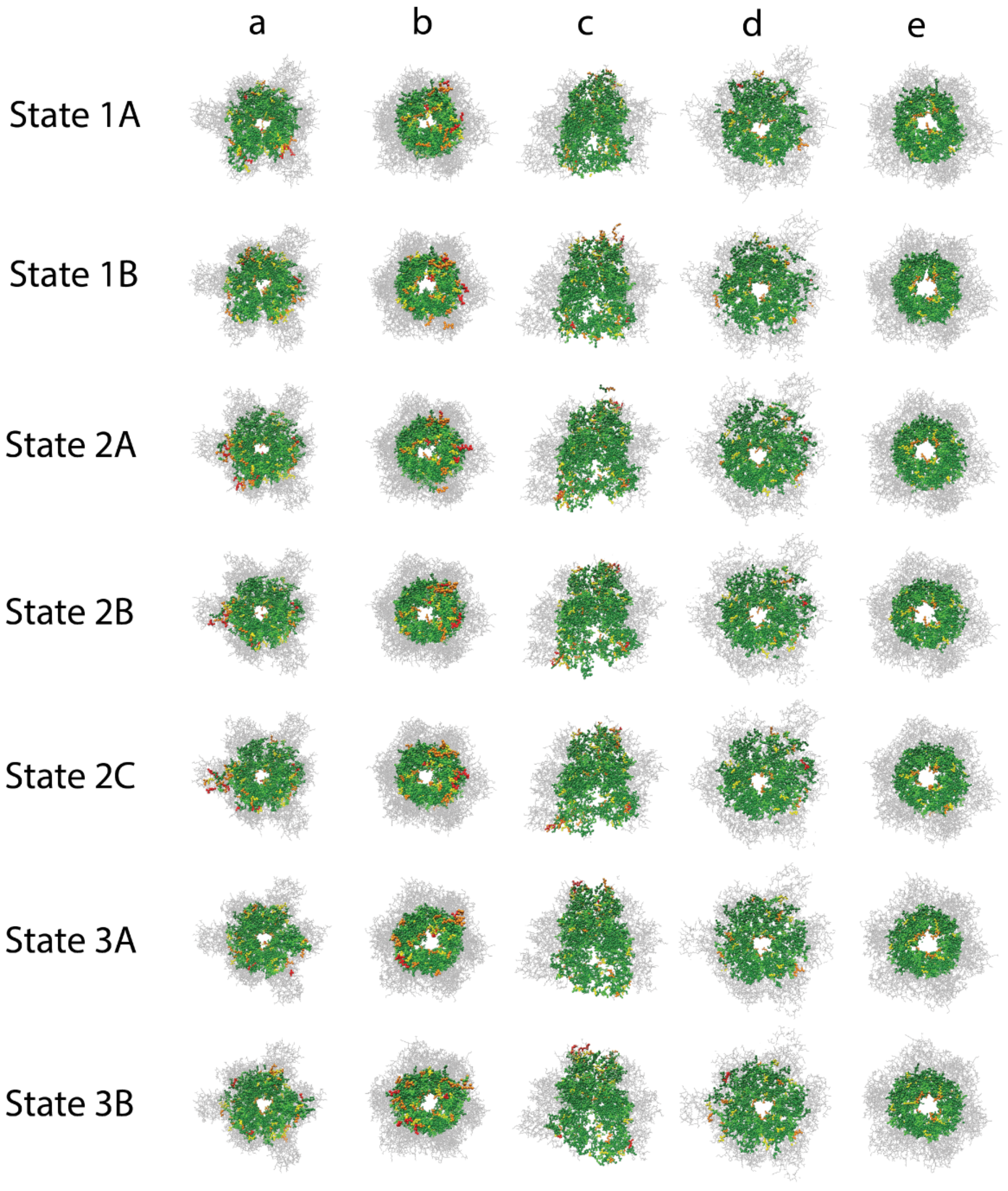


Figure 21: **Conservation profile of residues lining the inner cavity of F1 ATP synthase as detected by CICLOP.** The cavity is depicted as gray wire while each residue detected on the inner surface is colored according to the conservation score assigned to it by CICLOP. The highly conserved residues are depicted as dark green spheres, moderately conserved residues as green spheres, neutral as yellow, moderately unconserved as orange and highly unconserved as red spheres respectively. Panel a and b depict the bottom and top views of the cavity, c depicts a longitudinal section, while d and e depict a transverse section of the cavity as observed from the bottom and top respectively.

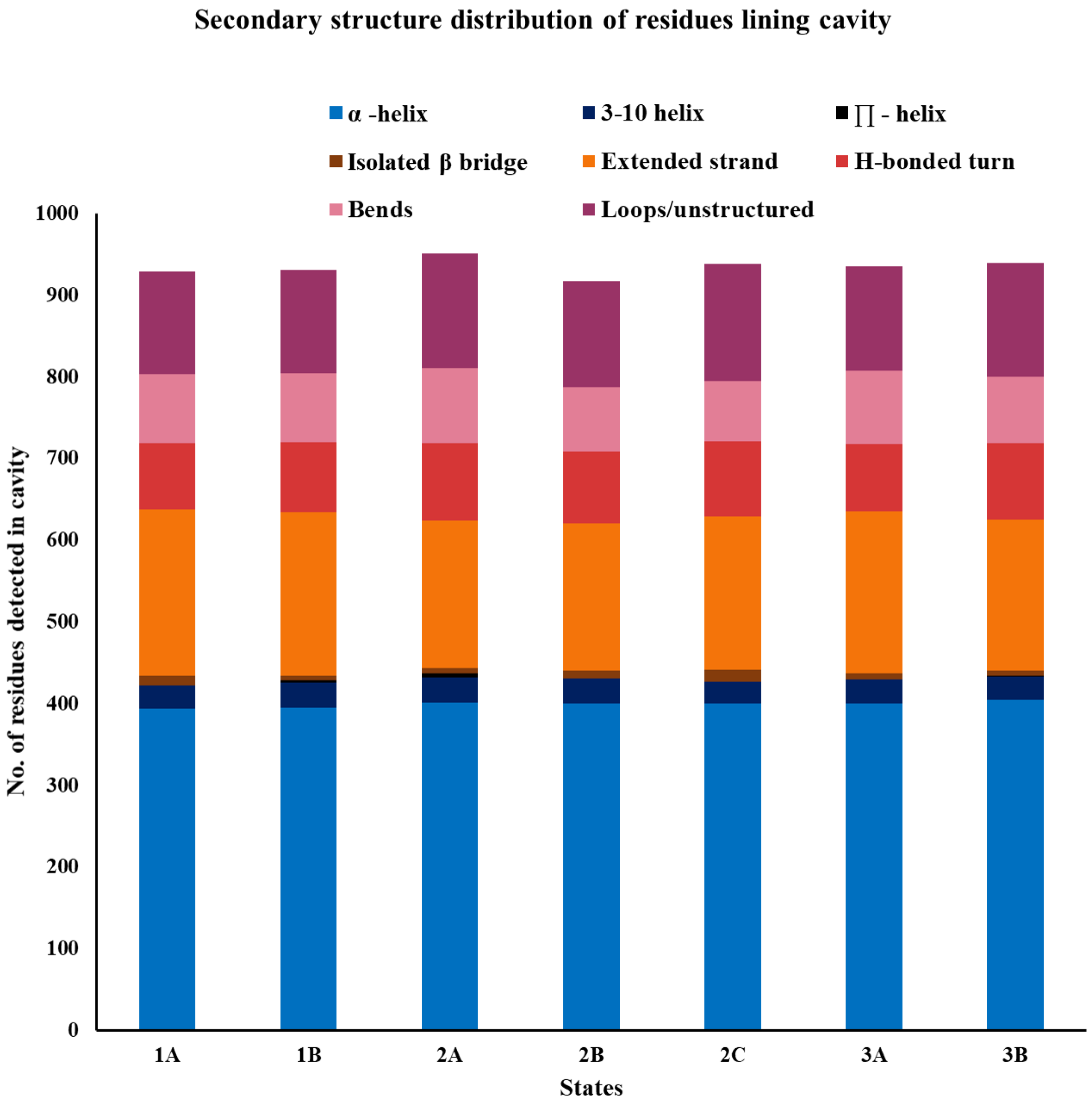


Figure 22: Secondary structure distribution of residues lining the cavity of ATP synthase F1 domain obtained using CICLOP. The number of residues participating in the various structural components is largely conserved in the different phases of the protein's rotation.

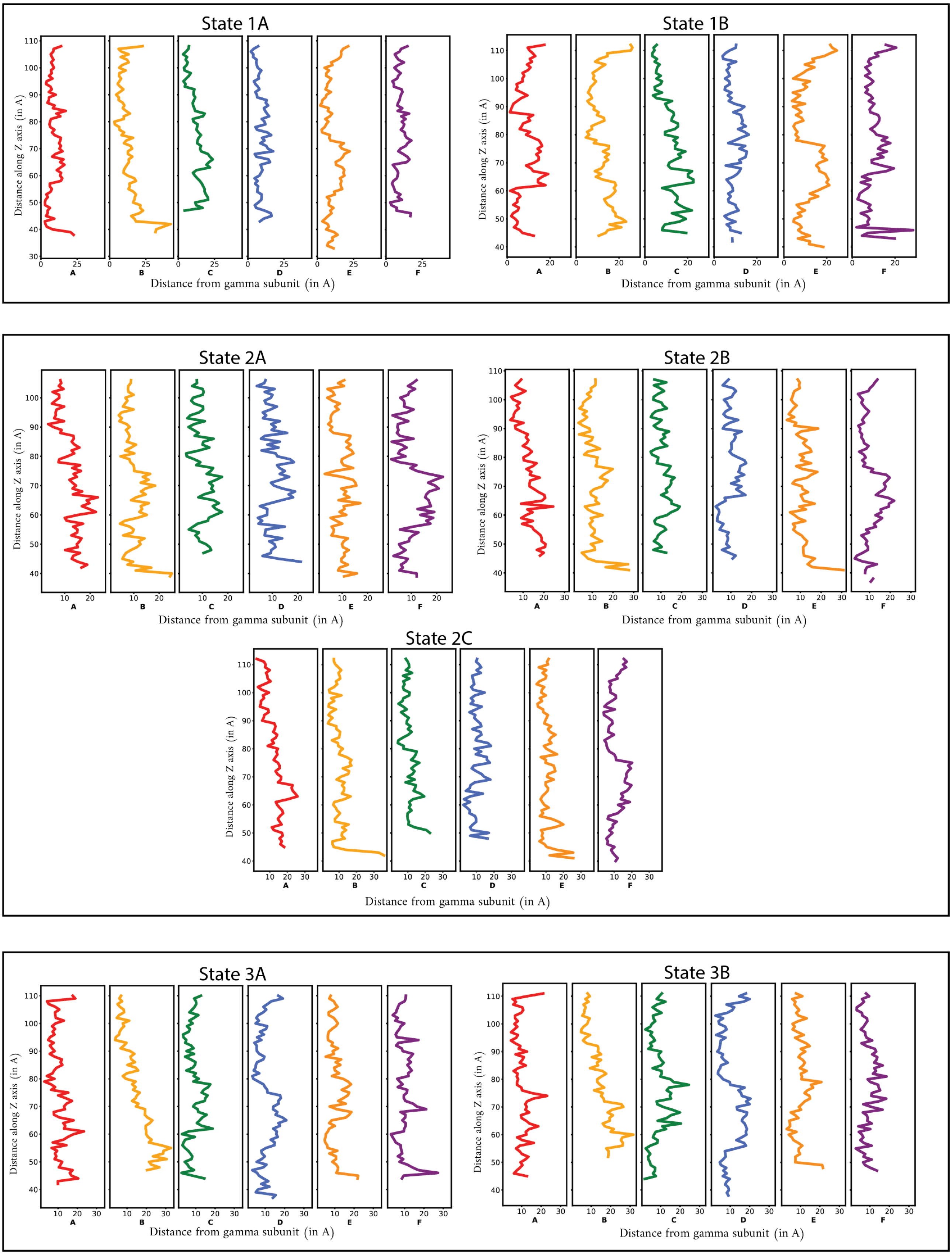


Figure 23: **Minimum distance from chain G to all the constituent chains of the cavity.** The figure quantifies the distance of $\gamma$ subunit (chain G) to all the chains constituting the cavity (chain A-F) respectively, highlighting the orientation as well as curvature of the rotor towards an $\alpha$ $\beta$ interface during the rotary cycle.

**Table 1: The PDB IDs used for the benchmarking study as well as the number of inner residues detected by each tool are indicated.**

| | PDBID | Total atoms | Total residues | Ciclop | Mole online | PoreWalker | Caverweb |
|---|---|---|---|---|---|---|---|
| 1 | **2MAL** | 643 | 93 | 54 | 121 | NaN | 101 |
| 2 | **2AWC** | 1232 | 130 | 76 | 145 | 64 | 12* |
| 3 | **5A64** | 3268 | 395 | 143 | 249 | 97 | 815 |
| 4 | **4EWS** | 4044 | 472 | 223 | 1129 | 192 | 831 |
| 5 | **3F6K** | 5600 | 661 | 253 | 366 | 151 | 47* |
| 6 | **6I5O** | 5723 | 560 | 226 | 208 | DNR | 30 |
| 7 | **6W5S** | 9700 | 1194 | 495 | 420 | 256 | 315 |
| 8 | **1EK9** | 11426 | 1284 | 667 | 3620 | 535 | 5672* |
| 9 | **6IA5** | 12022 | 1389 | 527 | 684 | DNR | 2162 |
| 10 | **6VAM** | 13360 | 1712 | 672 | 632 | 528 | 451 |
| 11 | **4NMD** | 16213 | 1959 | 735 | 1472 | 283 | DNR |
| 12 | **6V0H** | 16602 | 2178 | 807 | 24 | 311 | 6589 |
| 13 | **5DFA** | 18308 | 2054 | 712 | 498 | DNR | 279 |
| 14 | **4ADS** | 23152 | 2994 | 1015 | 762 | 104 | DNR |
| 15 | **5WC3** | 32490 | 4170 | 1536 | NaN | DNR | DNR |
| 16 | **1RSC** | 36846 | 4608 | 1450 | 58 | 466 | DNR |
| 17 | **6V0C** | 37920 | 4992 | 1686 | 3213 | 556 | DNR |
| 18 | **6XGR** | 38034 | 4950 | 1857 | NaN | 592 | DNR |
| 19 | **2WVW** | 42856 | 5416 | 1862 | 3311 | 290 | DNR |
| 20 | **4K0J** | 44018 | 5742 | 1955 | 6625 | DNR | DNR |
| 21 | **6VAI** | 45584 | 5874 | 2265 | 2143 | 1369 | DNR |
| 22 | **1AON** | 58870 | 8015 | 2707 | 1126 | 1154 | DNR |
| 23 | **5ZDH** | 66045 | 8715 | 3638 | 1392 | DNR | DNR |
| 24 | **3J9Q** | 99648 | 13224 | 4256 | 1034 | DNR | DNR |
| 25 | **3J9R** | 103824 | 13788 | 4121 | 3675 | DNR | DNR |
| 26 | **7CYC** | 302100 | 38580 | 899 | 7836 | DNR | DNR |

NaN : No tunnels detected

DNR : No output obtained. Did not run

[number]* : Ran manually. Starting point provided.
